# Neuromesodermal Progenitors Advance Network Formation of Spinal Neurons and Support Cells in Neural Ribbons *In Vitro* and Unprotected Survival in a Rat Subacute Contusion Model

**DOI:** 10.1101/2020.11.10.374876

**Authors:** Zachary T. Olmsted, Cinzia Stigliano, Annalisa Scimemi, Brandon Marzullo, Tatiana Wolfe, Jose Cibelli, Philip J. Horner, Janet L. Paluh

**Affiliations:** State University of New York Polytechnic Institute, Colleges of Nanoscale Science and Engineering, Nanobioscience Constellation, Albany, NY 12203; Houston Methodist Research Institute, Department of Neurosurgery, Center for Neuroregeneration, 6670 Bertner Ave. R10-North, Houston, TX 77030; State University of New York at Albany, Biological Sciences, 1400 Washington Avenue, Albany, NY, 12222; SUNY Buffalo Genomics and Bioinformatics Core, New York State Center of Excellence in Bioinformatics and Life Sciences, 701 Ellicott St. Buffalo, NY, 14222; Michigan State University, Dept. of Animal Science, College of Agriculture and Natural Resources and Large Animal Clinical Sciences, College of Veterinary Medicine, East Lansing, MI 48824

## Abstract

Improved human stem cell interventions to treat CNS trauma requires continued expansion of *in vitro* models and delivery platforms to fill gaps in analysis and treatment. Transplanted neural stem cells (NSCs) face unique, multi-faceted challenges beyond survival that include differentiation, maturation, and integration into a complex cytokine-releasing microenvironment that impinges on a multipotent cell type. Alternate strategies to transplant neurons and neuronal networks deserve reevaluation, particularly since novel differentiation protocols mimicking region-specific developmental and positional cues have recently emerged. To investigate transplantation of neurons and their early networks, we generate *in vitro* neural ribbons containing spinal neurons and support cells anatomically matched for cervical spinal cord injury (SCI). These glutamate-responsive, electrically-active neural ribbons apply a new hiPSC differentiation strategy transiting through neuromesodermal progenitors (NMps) to derive developmentally relevant spinal motor neurons (SMNs), interneurons (INs), and oligodendrocyte progenitor cells (OPCs). Bioinformatic profiling validates region-specific identities. Neurons and neuronal networks are functionally evaluated for action potential firing, calcium signaling, population activity, and synaptogenesis. NMp-derived neurons survive *in vivo* within the subacute phase hemi-contusion injury cavity when delivered either as free suspension or as encapsulated networks of pre-formed CNS cytoarchitectures. Delivery as encapsulated networks further supports survival of lower cell numbers and rapid graft penetration into host tissue. Neural network ribbons therefore provide a novel intermediary approach between cell suspensions and complex organoids for investigating network formation and early transplantation events with hiPSC-derived neurons, providing flexibility to rapidly tune cell type(s), cell ratios, and traceable biomarkers.

**Significance Statement:** In the two decades since human stem cell technologies have emerged, the challenge has remained to improve the developmentally relevant derivation of therapeutic cells. The ability to now generate anatomically matched neurons for SCI necessitates a re-evaluation of these cells and their networks *in vitro* and *in vivo*. In this study, we apply developmental cues via neuromesodermal progenitors to generate spinal neurons from hiPSCs. Genetic and functional evaluation of these cells as *in vitro* neuronal networks, due to their capacity to survive and graft effectively within the rat subacute contusion cavity, offer novel approaches for customizing SCI transplantation. This work demonstrates a strategy to develop transplantable, chemically-responsive networks linking *in vitro* models with injury customization towards improved *in vivo* outcomes.

## Introduction

Human neuronal precursors derived from stem cells have recently begun to show therapeutic efficacy in CNS disorders requiring cell replacement. At the forefront in the clinic is Parkinson’s disease (Barker et al., 2017), wherein neurodegeneration is initially restricted to a local population of dopaminergic neurons in the substantia nigra of the midbrain (DeMaagd and Philip, 2015). The homotypic approach to deliver developmentally patterned dopaminergic precursor cells in suspension to this region has been instrumental in enabling robust engraftment, *in vivo* neuronal maturation and circuit repair (Kikuchi et al., 2017; Xiong et al., 2020). In other neurological disorders incurred by acute insult, such as in stroke (Baker et al., 2019) or SCI (Silva et al., 2014), the spectrum of severity, anatomical distribution, and breadth of neural circuits affected both within a single subject and between subjects will likely preclude any single cell type-based therapy. As well, reproducible efficacy of NSC therapies to maximize functional improvement and clinical outcomes across these scenarios has remained at a preliminary stage (Katoh et al., 2019). Focused early integration studies continue to be important for achieving success and reproducibility in animal models that are particularly challenging to predict when using unprotected NSCs delivered in suspension. While some investigation into the transplantation of encapsulated cell types has been done (Tsintou et al., 2015; Ahuja et al., 2016; Liu et al., 2018), these studies primarily address multipotent cell protection and do not explore the transplantation of neurons, pre-configured neuronal networks in biomaterials, nor the ability to customize 3D neuronal networks for a variety of *in vitro* and *in vivo* applications.

The revelation of new details in spinal cord developmental principles and their implementation into stem cell research have combined to generate homotypic spinal neurons (Gouti et al., 2014; Sagner and Briscoe, 2019). Here we expand such approaches through anatomically and physiologically relevant neuromesodermal progenitor (NMp) and spinal cord NSC (scNSC) intermediates. This approach avoids the use of default NSCs with forebrain identity that have been applied in many published SCI studies, allowing the generation of developmentally appropriate spinal motor neurons (SMNs) along with a subpopulation of co-developed interneurons (INs) and oligodendrocyte progenitor cells (OPCs). By patterning spinal neurons and support cells within neural ribbon constructs, we demonstrate immature network formation with output of stereotypic firing activity in response to glutamate stimulation. *In vitro*, the maturing neurons satisfy a suite of functional and electrophysiological hallmarks, which we parallel with whole-transcriptome time course analysis during differentiation. This study is the first to fully characterize NMp-differentiated SMNs transcriptionally, functionally, and at the protein level in culture, as networked ribbons, and *in vivo*. By MRI of neural ribbons containing SMNs labeled with superparamagnetic iron oxide nanoparticles, we demonstrate graft linear placement and retention in the healthy rat spinal cord. Using a cervical hemi-contusion injury model in rat, we show that our homotypic SMNs survive transplantation when delivered either as suspensions or as SMN-encapsulated neural ribbons. Biomarker analysis of spinal cord sections reveals robust survival of the homotypic cervically-matched neurons eight days after transplantation, and rapid engraftment within the host tissue at the border of the subacute injury cavity. This study demonstrates that homotypic, maturing spinal neurons derived through NMps survive the subacute phase SCI microenvironment. Our novel neural ribbon platform extends capabilities for design of pre-configured functioning networks for *in vitro* modeling or CNS delivery analysis.

## Materials and Methods

### Human hiPSC maintenance and neuronal differentiation

The African American hiPSC line F3.5.2 was developed in our laboratory (Chang et al., 2015) and is extensively characterized (Tomov et al., 2016; Olmsted et al. 2020). F3.5.2 was cultured in mTeSR Plus (STEMCELL Technologies) supplemented with 1x penicillin-streptomycin (P-S; Gibco) on hESC-qualified Corning Matrigel (1:100) at 37°C, 5% CO_2_. hiPSC colonies were passaged 1:6 every 4 to 7 days with Gentle Cell Dissociation Reagent (STEMCELL Technologies). Genomic integrity was validated by G-band karyotyping (Cell Line Genetics, Madison, WI). On day 0 of differentiation, hiPSC colonies at 60-70% confluency were rinsed in DMEM/F-12 medium and Neuromesodermal Progenitor Medium (NMPM) was added. N2B27 basal medium: 1:1 DMEM/F-12:Neurobasal Plus medium, 2% (vol/vol) B-27 Plus supplement, 1% (vol/vol) N-2 supplement, 1x GlutaMAX, 1x MEM Non-Essential Amino Acids, 1x P-S (Gibco); NMPM: N2B27 supplemented with 40 ng/ml recombinant human (rh) FGF2 (R&D Systems), 40 ng/ml rhFGF8 (R&D Systems), 2 μM CHIR99021 (Tocris Bioscience), 10 μM DAPT (Millipore Sigma), 10 μM SB431542 (Tocris Bioscience), 100 nM LDN193189 (Tocris Bioscience), 0.36 U/ml heparin (Millipore Sigma). NMPM was changed daily. At day 5, NMps were passaged with Gentle Cell Dissociation Reagent as 50-100 μm aggregates in cervical patterning medium (CPM). CPM: N2B27 supplemented with 100 nM Retinoic Acid (RA; Millipore Sigma), 200 nM Hh-Ag1.5. 10 μM Y-27632 (Tocris Bioscience) was added during passages and replaced with fresh medium. scNSC cultures were passaged 1:3 in 12-well plates and maintained at 90-100% confluency for 2-3 days before subsequent high-density passages (1:2). Neural cells were cultured on Matrigel for the entirety of differentiation. scNSCs were patterned in CPM to day 25 and transitioned to terminal differentiation medium (TDM). TDM: N2B27 supplemented with 10 ng/ml rhBDNF, 10 ng/ml rhGDNF, 1 μM dibutyryl cyclic-AMP (dbcAMP; Millipore Sigma). For culture purification during motor neuron progenitor (MNP) and SMN differentiation, 24 - 48 h neurosphere (NS) cultures were formed in 6-well plates (CELLTREAT) using a Scilogex MX-M microplate mixer at ~120 rpm and freshly seeded. The differentiation protocol was repeated >60 times over the course of this study.

### Lentivirus transduction

GFP labeling of cells using lentivirus transduction was performed similarly to described (Taylor et al., 2006) at the hiPSC pluripotent stage. In brief, hiPSCs were seeded onto freshly coated 6-well plates 3 days prior to transduction with premade LV-CAG-eGFP lentivirus (Kerafast FCT149, 1 × 10^8^ CFU/ml). Polybrene (2 μg/ml; Millipore Sigma) was added to 2 ml mTeSR Plus. hiPSCs were transduced at 5x magnification of infection by centrifugation with the mTeSR-PB-LV medium for 1 h at 32°C, 1,200 × *g* and subsequent incubation at 37°C, 5% CO_2_ for 18 h. Cultures were rinsed 2x with DMEM/F-12 and cultured in mTeSR Plus for 48 h prior to visual inspection by fluorescence microscopy. Transduced hiPSCs were selected and expanded in the presence of 2 μM puromycin and cryo-stored in mFreSR (STEMCELL Technologies) or further differentiated to OPCs as described below.

### OPC differentiation, sorting, and SMN co-culture

Similarly to SMNs, OPCs were patterned through NMp and scNSCs stages, but excluding DAPT (Goldman and Kuypers, 2015). For neural ribbon encapsulation, LV-CAG-eGFP transduced hiPSCs were used as starting material. At day 19, 24 h short-term NS suspension culture was used as a purification strategy. NS were seeded at day 20 in OPC medium (OPCM) (Khazaei et al., 2017). OPCM: N2B27 (no Vitamin A) supplemented with 10 ng/ml rhIGF-1 (R&D Systems), 20 ng/ml rhFGF2 (R&D Systems), 20 ng/ml rhPDGF-AA (R&D Systems), 60 ng/ml tri-iodothyronine (T3; Millipore Sigma), 0.36 U/ml heparin (Millipore Sigma). At day 36, cultures were transitioned to OPC maturation medium (OPMM). OPMM: 10 ng/ml rhIGF-1, 25 μg/ml insulin, 1 μM dbcAMP, 20 μg/ml ascorbic acid, 60 ng/ml T3.

For co-culture with SMNs, GFP-OPCs at day 35 were sorted manually using the MidiMACS kit (Miltenyi Biotec 130-090-312) with LS columns and Anti-PE MicroBeads (Miltenyi Biotec 130-048-801) according to manufacturer’s instructions. Cells were labeled with IgM PE-O4 antibody (R&D Systems FAB1326P) at 10 μl/10^6^ cells. GFP-OPCs were retained and mixed 1:5 with SMNs. Co-cultures were seeded and maintained for 1 w in modified N2B27 medium, deemed bundling medium, prior to neural ribbon co-encapsulation. Bundling Medium: N2B27 supplemented with 10 ng/ml rhBDNF, 10 ng/ml rhGDNF, 10 ng/ml rhNGF, 20 ng/ml rhPDGF-AA, 10 ng/ml rhIGF-1, 60 ng/ml T3, 100 μM dbcAMP, 25 μg/ml insulin, 20 μg/ml ascorbic acid.

### Phase contrast imaging and immunofluorescence

Phase contrast images were acquired using a Nikon Eclipse TS100 microscope (10x/0.25 Ph1 ADL and 20x/0.40 Ph1 ADL objectives; Qimaging Retiga 2000R camera) or Zeiss Invertoskop 40C (5x/0.12 CP-Apochromat, 10x/0.25 Ph1 A-Plan and 20x/0.30 Ph1 LD A-Plan objectives; Olympus DP22 color camera). For immunofluorescence (IF) imaging, cells were seeded into Matrigel-coated glass-bottom chambers (Lab-Tek II 4-chambered cover glass; Nunc, %155382) and fixed with 10% buffered formalin for 30 min at specified time points. Samples were permeabilized for 5 min in 0.1% Triton-X-100 and blocked for 30 min in 1% BSA fraction V (1x PBS). Primary antibodies were applied in 1 ml fresh blocking buffer and incubated at 4°C overnight (**Table 1**). Samples were rinsed thoroughly in 1x PBS before applying immunoglobulin- and species-matched AlexaFluor secondary antibodies (Invitrogen) for 1 h in the dark with DAPI (4°C). Cells were imaged directly in chambered cover glass wells or mounted with ProLong Gold Antifade (Invitrogen). Fluorescence microscopy was performed using a Zeiss Axio Observer.Z1 inverted fluorescence microscope (20x/0.8 air and 63x/1.4 oil Plan-Apochromat DIC objectives). Images were acquired using an Hamamatsu ORCA ER CCD camera and Zeiss AxiovisionRel software (ver. 4.8.2). For adherent cultures, 6- to 30-slice Z-stacks were gathered at 1 μm separation distance and compressed using the Extended Focus feature. Uniform exposure times were maintained across samples for identical antibodies. If necessary, images were adjusted linearly for brightness in Keynote or ImageJ. A Leica confocal TCS SP5 II system was used for lower magnification images (10x/0.30 HCX PL FLUOTAR air objective lens), acquired using Leica Application Suite Advanced Fluorescence software.

**Table 1.** Antibodies Used *In Vitro*.

| Target | Company; Cat# | Species; Clone | Dilution |
| --- | --- | --- | --- |
| α-tubulin | Invitrogen; 62204 | Mouse mAb IgG<sub>1</sub>; DM1A | 1:1,000 |
| SOX2 | R&D Systems; MAB2018 | Mouse mAb IgG<sub>2A</sub>; 245610 | 1:1,000 |
| SOX2 | R&D Systems; AF2018 | Goat pAb IgG | 1:1,000 |
| SOX1 | R&D Systems; AF3369 | Goat pAb IgG | 1:500 |
| Nestin | R&D Systems; MAB1259 | Mouse mAb IgG<sub>1</sub>; 196908 | 1:1,000 |
| Pan-Cadherin | Abcam; ab6528 | Mouse mAb IgG<sub>1</sub>; CH-19 | 1:1,000 |
| NCAM-1 | R&D Systems; AF2408 | Goat pAb IgG | 1:1,000 |
| NG2 (*CSPG4*) | eBioscience; 14-6504-80 | Mouse mAb IgG<sub>2A</sub>; 9.2.27 | 1:1,000 |
| CDX-2 | R&D Systems; MAB3665 | Mouse mAb IgG<sub>1</sub>; 963809 | 1:1,000 |
| Hox-C6 | Santa Cruz Biotechnologies;<br>sc-376330 | Mouse mAb IgG<sub>2A</sub>; B-7 | 1:500 |
| PAX6 | DSHB; PAX6 | Mouse mAb IgG<sub>1</sub>; PAX6 | 1:500 |
| Nkx-6.1 | DSHB; F55A12 | Mouse mAb IgG<sub>1</sub>; F55A12 | 1:250 |
| OLIG2 | R&D Systems; AF2418 | Goat pAb IgG | 1:500 |
| TUJ1 (βIII-tubulin,<br>TUBB3) | BioLegend; 802001 | Rabbit pAb IgG;<br>Poly18020 | 1:2,000 |
| MAP2 | Abcam; ab32454 | Rabbit pAb IgG | 1:1,000 |
| SMI312 | BioLegend; 837904 | Mouse mAb IgG<sub>K</sub>/IgM<sub>K</sub><br>cocktail; SMI312 | 1:1,000 |
| SMI311R | BioLegend; 837801 | Mouse mAb IgG/IgM<br>cocktail; SMI311R | 1:1,000 |
| NeuN | Millipore; MAB377 | Mouse mAb IgG<sub>1</sub>; A60 | 1:1,000 |
| ISL-1&2 | DSHB; 39.4D5 | Mouse mAb IgG<sub>2B</sub>; 39.4D5 | 1:500 |
| ISL1 | Millipore Sigma;<br>HPA057416 | Rabbit pAb IgG | 1:1,000 |
| HB9 (*MNR2/MNX1*) | DSHB; 81.5C10 | Mouse mAb IgG<sub>1K</sub>;<br>81.5C10 | 1:100 |
| ChAT | R&D Systems; MAB3447 | Mouse mAb IgG<sub>1</sub>;<br>#334008 | 1:1,000 |
| LHX3 | Santa Cruz Biotechnologies;<br>sc-293411 | Mouse mAb IgG<sub>2A</sub>; 2C10 | 1:1,000 |
| FOXP1 | Santa Cruz Biotechnologies;<br>sc-398811 | Mouse mAb IgG<sub>1</sub>; A-2 | 1:1,000 |
| RALDH2 (*ALDH1A2*) | Millipore Sigma; ABN420 | Rabbit pAb IgG | 1:1,000 |
| Peripherin | Santa Cruz Biotechnologies; sc-377093 | Mouse mAb IgG<sub>2A</sub>; A-3 | 1:1,000 |
| AnkG | Antibodies Inc.; 75-147 | Mouse mAb IgG<sub>2B</sub>; N106/65 | 1:200 |
| β4-Spectrin | Antibodies Inc.; 73-376 | Mouse mAb IgG<sub>1</sub>; N393/2 | 1:200 |
| Kv1.2 (*KCNA2*) | Novus Biologicals; NBP1-42802SS | Rabbit pAb IgG | 1:1,000 |
| Synapsin 1 (*SYN*) | Millipore Sigma; AB1543 | Rabbit pAb IgG (serum) | 1:1,000 |
| GluR1 (*GRIA1*) | Antibodies Inc.; 75-327 | Mouse mAb IgG<sub>1</sub>; N355/1 | 1:1,000 |
| GluN2A (*GRIN2A*) | Antibodies Inc.; 75-288 | Mouse mAb IgG<sub>2A</sub>; N355/1 | 1:1,000 |
| PSD-95 (*DLG4*) | Abcam; ab18258 | Rabbit pAb IgG | 1:500 |
| GAP-43 | Encor; MCA-5E8 | Mouse mAb IgG<sub>1</sub>; GAP43 | 1:1,000 |
| STMN2 | Novus Biologicals; NBP149461 | Rabbit pAb IgG | 1:1,000 |
| Nkx-2.2 | DSHB; 74.5A5 | Mouse mAb IgG<sub>2B</sub>; 74.5A5 | 1:500 |
| O4 | R&D Systems; MAB1326 | Mouse mAb IgM; #O4 | 1:500 |
| O4 | Chemicon; MAB345 | Mouse mAb IgM; 81 | 1:500 |
| PE-O4 | R&D Systems; FAB1326P | Mouse mAb IgM; #O4 | MACS |
| GFAP | R&D Systems; MAB2594 | Mouse mAb IgG<sub>1</sub>; #273807 | 1:1,000 |
| CHX10 | Santa Cruz Biotechnologies; sc-374151 | Mouse mAb IgG<sub>2B</sub>; D-11 | 1:1,000 |
| GATA-3 | R&D Systems; MAB6330 | Mouse mAb IgG<sub>2B</sub>; 634913 | 1:1,000 |
| PAX2 | Invitrogen; 71-600-0 | Rabbit pAb IgG | 1:1,000 |
| LBX1 | Invitrogen; PIPA564884 | Rabbit pAb IgG | 1:500 |
| TLX3 | Abcam; ab184011 | Rabbit pAb IgG | 1:1,000 |
| LHX9 | Abcam; ab224357 | Rabbit pAb IgG | 1:1,000 |
| BRN3A | Millipore Sigma; AB5945 | Rabbit pAb IgG | 1:1,000 |
| GFP | Invitrogen; A-6455 | Rabbit pAb IgG | 1:1,000 |
| Desmin | R&D Systems; AF3844 | Goat pAb IgG | 1:1,000 |
| Myogenin | R&D Systems; MAB6686 | Mouse mAb IgG<sub>1</sub>; #671038 | 1:500 |
| Myosin Heavy Chain | R&D Systems; MAB4470 | Mouse mAb IgG<sub>2B</sub>; #MF20 | 1:1,000 |

### Calcium imaging

SMN cultures were incubated at 37°C with 5 μM Fluo-4 AM (Invitrogen) for 30 min in Neurobasal A medium (no supplements). Cultures were then recovered in fresh TDM for 30 min and imaged in HBSS using the Zeiss Axio Observer.Z1 system with the 20x air objective described above. Time lapse series were captured at an exposure time of 50 ms obtained in 200 ms intervals for a total duration of 1.5 or 2.5 min. ΔF/F over time was determined using PhysImage software (ImageJ distribution) as previously described (Hayes et al., 2018; Faustino Martins et al., 2020).

### Neuromuscular junctions

NMJ assays were performed similarly to as described in previous studies (Harper et al., 2004; Miles et al., 2004). Briefly, rat L6 myoblasts (ATCC CRL-1458) were seeded into Matrigel-coated Lab-Tek II 4-chambered cover glass and grown to confluency in DMEM/F-12, 10% FBS, 1x P-S medium. At 100% confluency, myoblasts were cultured in N2B27 medium supplemented with 100 nM RA, 200 nM Hh-Ag1.5 to drive multinucleated skeletal myotube formation. 24 h NS (day 38) were seeded onto myotubes at 5 - 8 NS per well in TDM and cultured for four days prior to fixation and fluorescence microscopy (day 42 SMNs).

### Neural ribbon encapsulation and dissolution

Lyophilized ultrapure sodium alginate (DuPont Novamatrix) was resuspended in sterile 0.9% NaCl (w/w) to 1.5% (w/w) and stored at 4°C. Two alginate products were used that are ProNova SLM100 (MWT: 150-250 kDa; G/M ratio < 1; viscosity 170 mPa*s; endotoxin < 25 EU/g) or Novatach LVM GRGDSP-coupled alginate (G/M ratio < 1, endotoxin < 10; peptide 0.015 μmol/mg). Peptide-conjugated alginate was used for *in vitro* assays while unconjugated alginate was used for *in vivo* grafting studies. 5 mg/ml PureCol EZ gel Type 1 Collagen solution (Advanced Biomatrix #5074) was included 1:3 with 1.5% alginate to provide initial ECM in neural ribbons prior to extrusion. ProNova-Collagen and RGD-Alginate-Collagen hydrogels are referred to here as NovaCol and RGD-Col, respectively. Differentiating cultures were lifted as aggregates using Accutase and distributed to 1.5 ml Eppendorf tubes in DMEM/F-12. A subset of aggregates was dissociated to single cells and counted. Cells were centrifuged at 350 × g for 5 min and the supernatant was aspirated. Cell pellets were combined first with non-gelled PureCol solution and resuspended gently at room temperature. 1.5% alginate solution was then added slowly while mixing, avoiding bubbles, to a final cell suspension concentration equivalent of 1× 10^8^ cells/ml. 40 μl of this suspension was loaded directly into a luer lock steel tip (34G, inner diameter 60 μm, 32G, inner diameter 100 μm, or 30G, inner diameter 150 μm, Nordson EFD) and attached to a sterile 3 ml BD syringe. Contents were dispensed directly into 100 mM CaCl_2_ crosslinking solution normal to the liquid surface for immediate templated formation of 60 μm neural ribbons. CaCl_2_ was aspirated completely after 1 min and the neural ribbons were resuspended in cell culture medium. Suspended neural ribbons were subsequently incubated (37°C, 5% CO_2_) to drive interpenetrating network formation by collagen gelation. Ribbon quality was monitored by visual inspection. Cell suspensions were excluded when forming empty hydrogel ribbons. Neural ribbons were either used immediately for downstream experiments or retained in culture. To collect cells for RNA extraction (RNA-Seq) or cover glass seeding (viability, electrophysiology), alginate was rapidly dissolved in 1.6% sodium citrate solution and cells were pelleted by centrifugation.

### Neural ribbon immobilization and imaging

To template and image neural ribbon cytoarchitecture, RGD-Col ribbons were formed as described above directly into the 12 mm viewing area of Nunc 35 mm glass bottom dishes (ThermoFisher #150680). CaCl_2_ was aspirated after crosslinking and the ribbons were manually aligned. To immobilize ribbons in position for imaging assays, 100 μl of PureCol EZ gel solution (5 mg/ml in DMEM) was gently applied over the ribbons, filling the viewing area, and incubated for 45 min at 37°C. After gelation, 1.5 ml of culture medium was added. To constrain neurite outgrowth to within the neural ribbon body, 4 μg/ml Aggrecan containing inhibitory CSPGs was included in the collagen embedding matrix prior to gelation. This cytoarchitecture was further promoted by culturing in bundling medium, described above. Cultures were maintained for 7 - 12 days in this platform before collagen gel began to degrade. The exclusion of Aggrecan enables neurite extension orthogonally outward from the ribbon body in addition to the interior. MitoTracker Green FM dye (Invitrogen) was added directly to culture medium as per manufacturer’s instructions and diffused into the collagen gel for 1 hr prior to imaging in HBSS without phenol red. For IF experiments, collagen solution was diluted 1:1 with DMEM/F-12 to enhance antibody permeability and wash steps were extended to 15 min each at room temperature. The IF protocol was modified for neural ribbons and was performed in 25 mM PIPES/10 mM HEPES/2.5 mM CaCl_2_ buffer to prevent dissolution by high phosphate concentration and Ca^2+^ leaching.

### Microelectrode array electrophysiology and analyses in adherent cultures

To monitor spontaneous neuronal activity in adherent SMN cultures, we seeded MNP-NS or neural ribbon aggregates directly onto 60MEA200/30iR-Ti microelectrode arrays (MEA; Multi Channel Systems). MEA chips contained single wells with 60 electrodes per well, each of 30 μm diameter with 200 μm inter-electrode spacing (input impedance 30-50 kΩ). One electrode per MEA chip served as a reference electrode. To obtain recordings, we used an MEA2100 head stage system connected to an MCS-IFB-in-vitro interface board. Temperature was maintained at 37°C during recordings using a temperature controller. Arrays were coated with Poly-D-Lysine and 20 μg/ml Laminin. On day 28 of differentiation, 12-15 NS were seeded in TDM and allowed to adhere for 2 days, at which point one-half medium changes were made (1 ml total volume). One-half medium changes were subsequently made twice per week. Recordings were acquired after NS culturing for 6 days with Multi Channel Experimenter software (v2.14.0) sampled at 25 kHz for at least 5 min. Spike threshold for detection was automatically set to 5x above standard deviation over background noise (1 ms pre trigger, 2 ms post trigger). Burst detection parameters were set as follows: 50 ms interval to start and end burst, 100 ms minimum inter-burst interval, 50 ms minimum burst duration, 4 minimum spikes per burst. Spike and burst analyses were performed using Multi Channel Analyzer (v2.14.0) in conjunction with NeuroExplorer (v5.022), and are shown for days 35 and days 52 in differentiation in accordance with patch clamp experimental time points. For glutamate stimulation, we prepared a 10 mM solution in N2B27 medium and added glutamate to a final concentration of 50 μM. Recordings were taken for a minimum of 5 min and up to 14 min. N = 4-5 separate MEA chips were used for quantification per time point. Generation of raw data traces and statistical analyses were performed using GraphPad Prism 8 in conjunction with Microsoft Excel.

### Microelectrode array electrophysiology recordings from neural ribbons

To investigate spiking, bursts, and networks bursts in 3D neural ribbons, we used 60HexaMEA40/10 single-well MEA chips. Each hexagonal array contained 60 electrodes, each of 10 μm diameter with 40 μm inter-electrode spacing that is suitable to neural ribbon geometry. Neural ribbon recordings were taken as acute measurements using day 35 SMNs encapsulated at day 28. To ensure electrode contact, neural ribbons were immobilized under a 12 mm coverslip compressed by a 5 g stainless steel weight. N = 3 neural ribbons were recorded under spontaneous conditions and after addition of 50 μM glutamate for a minimum of 6 min using the settings described above. For statistical analysis on the day of recording, glutamate addition and washout was performed three times for the same neural ribbon wherein recordings were separated by 30 min each to account for inactivation of glutamate ionotropic channels. Burst detection parameters were identical to adherent recordings described above. For network burst analysis, detection was set to minimum active channels of 4 and minimum simultaneous channels of 4, due to incomplete electrode coverage by the neural ribbons.

### Patch clamp electrophysiology

hiPSCs were differentiated as described to day 23, at which point adherent cultures were passaged using Gentle Cell Dissociation Reagent as aggregates and encapsulated in neural ribbons (1 × 10^8^ cells/ml equivalent) or as neurospheres in suspension culture. Neural ribbon and NS suspension cultures were maintained for 48 h. On day 25, NS were plated onto 18 mm Matrigel-coated cover glass (#1.5 thickness, Warner Instruments). Neural ribbons were dissolved in 1.6% sodium citrate and seeded at an equivalent cell density. We further compared neurons cultured in patterning versus maturation medium using 5 μl/ml BDNF and 5 μl/ml GDNF slow release PLGA microbeads (StemCultures; 5 ng/ml steady-state concentrations) at two time points, for a total of six conditions (**Table 2**). The experiments were performed on multiple differentiations per condition (N = 3–4). We added 10 ng/ml soluble rhBDNF and rhGDNF (R&D Systems) to the culturing medium on days of TDM media changes. For patch clamp studies, motor neurons were visually identified based on trapezoidal shape with more than two dendritic processes and the presence of high-contrast deposition in the cell cytoplasm (Sareen et al., 2013). The SMNs were transferred to a recording chamber perfused at 2.5 ml/min with a solution saturated with 95% O2/5% CO2 and containing (in mM): 119 NaCl, 2.5 KCl, 1.2 CaCl_2_, 1 MgCl_2_, 26.2 NaHCO_3_, 1 NaH_2_PO_4_, 22 glucose; 300 mOsm; pH 7.4. Whole-cell patch clamp recordings were made with a pipette solution containing (in mM): 120 KCH_3_SO_3_, 10 EGTA, 20 HEPES, 2 MgATP, 0.2 NaGTP, 290 mOsm, pH 7.2. The

**Table 2.** Electrophysiology Current Clamp Samples.

| Days in differentiation | Medium | PLGA neurotrophic factor beads | Neural ribbon (Y/N) | Cells patched (n = ) | Differentiations (N = ) |
| --- | --- | --- | --- | --- | --- |
| 33-35 | CPM<sup>a</sup> | None | No | 10 | 3 |
| 32-34 | TDM<sup>b</sup> | 5 µl/ml BDNF, GDNF | No | 9 | 3 |
| 32-34 | TDM | 5 µl/ml BDNF, GDNF | Yes | 13 | 3 |
| 45-51 | CPM | None | No | 11 | 3 |
| 51-52 | TDM | 5 µl/ml BDNF, GDNF | No | 11 | 4 |
| 51-52 | TDM | 5 µl/ml BDNF, GDNF | Yes | 13 | 3 |

<sup>a</sup>Cervical Patterning Medium (N2B27 + RA, Hh-Ag1.5)
<sup>b</sup>Terminal Differentiation Medium (N2B27 + BDNF, GDNF, dbcAMP)

resistance of the recording electrode was ~5 MΩ and was monitored throughout the experiments by applying a −50 pA, 10 ms pulse. Data were discarded if the input resistance changed more than 20% during the course of the experiment. All recordings were obtained under current-clamp configuration using a Multiclamp 700B amplifier and a 10 kHz low-pass filter (Molecular Devices). All traces were digitized at 10 kHz. Each cell was subject to a series of 5 ms (short) and 500 ms (long) current step injections. The amplitude of the short current step ranged between 50 and 350 pA. The amplitude of the long current step ranged between −100 and 350 pA. The amplitude of the short and long current injections were changed in steps of 10 pA. Each protocol was applied for 10 consecutive times. The data analysis was performed off-line with custom-made software (A.S.) written in IgorPro 6.36 (Wavemetrics). All experiments were performed at room temperature.

### Bulk RNA-Seq

Detailed information of RNA-Seq biological samples is provided in **Table 3**. SMNs were differentiated as described. RNA was extracted using the PureLink RNA Mini Kit (Invitrogen) according to manufacturer’s instructions. Neural cells from three wells of a 12-well plate were pooled for each sample for library preparation, and RNA-Seq was performed at the SUNY Buffalo Genomics and Bioinformatics Core. Per-cycle basecall (BCL) files generated by the Illumina NextSeq were converted to per-read FASTQ files using bcl2fastq version 2.20.0.422 using default parameters and no lane splitting. The quality of the sequencing was reviewed using FastQC version 0.11.5 and FastqScreen version 0.11.1. Quality reports were summarized using MultiQC version 1.7 (Ewels et al., 2016). No adapter sequences were detected so no trimming was performed. Genomic alignments were performed using hisat2 version 2.1.0 using default parameters (Kim et al., 2015). The Illumina provided UCSC hg38 from their igenomes database was used for the reference genome and gene annotation set. Sequence alignments were compressed and sorted into binary alignment map (BAM) files using samtools version 1.7. Counting of mapped reads for genomic features was performed using Subread featureCounts version 1.6.2 using the parameters -p -s 2 -g gene_name -t exon -B -C -Q 60 the annotation file specified with -a was from UCSC hg38 (Liao et al., 2014). Alignment statistics and feature assignment statistics were again summarized using MultiQC. The unique alignment rate of the paired-end reads for all samples was around 85%. The number of mapped reads assigned to features ranged from about 8 million to 16 million. Differentially expressed genes were detected using the Bioconductor package DESeq2 version 1.24.0. DESeq2 tests for differential expression using a negative binomial generalized linear models, dispersion estimates, and logarithmic fold changes (Love et al., 2014). DESeq2 calculates log2 fold changes and Wald test p-values as well as preforming independent filtering and adjusts for multiple testing using the Benjamini-Hochberg procedure to control the false discovery rate (FDR). Histograms were generated using Microsoft Excel and GraphPad Prism.

**Table 3.** RNA-Seq Samples.

| Day in differentiation | Stage | Purification | Differentiation | Wells pooled (12w plate) | Comments |
| --- | --- | --- | --- | --- | --- |
| 10 | scNSC rosettes | N/A | A | 3 | Biol. replicate |
| 10 | scNSC rosettes | N/A | B | 3 | Biol. replicate |
| 19 | MNP | N/A | A | 3 | CPM<sup>a</sup> at d5 |
| 25 | MNP | N/A | A | 3 | CPM at d5 |
| 28 | SMN | N/A | A | 3 | TDM<sup>b</sup> at d25 |
| 32 | SMN | NS (d28) | A | 3 | Plated d29 |
| 37 | SMN | NS (d28) | A | 3 | Plated d29 |
| 42 | SMN | NS (d28) | A | 3 | Plated d29 |

<sup>a</sup>Cervical Patterning Medium (N2B27 + RA, Hh-Ag1.5)
<sup>b</sup>Terminal Differentiation Medium (N2B27 + BDNF, GDNF, dbcAMP)

### Neural ribbon long-distance shipping and receiving

Neural ribbons were shipped overnight from SUNY Polytechnic (Albany, NY) to Houston Methodist (Houston, TX). Day 30 spinal neuron cultures were encapsulated in neural ribbons in 12-well plates, resuspended in TDM supplemented with 10 μM ROCK inhibitor, and incubated at 37°C for 1 h. Ribbon suspensions were then transferred to 1.8 ml cryovials using 1,000-μl pipette tips and immediately shipped on ice (4°C) via FedEx (20-22 h total in transit). At the destination laboratory, neural ribbons were recovered in a 37°C humidified cell culture incubator prior to the day of transplantation in rat.

### SPIO labeling and collagen phantom injections

At day 29, confluent neuronal cultures were incubated with FeraTrack MRI SPIO nanoparticles (Miltenyi Biotec) overnight in 12-well plates. Cell viability versus SPIO concentration analysis was performed after 24 h incubation. ~100 μg Fe / 1 × 10^6^ cells (40 μl Miltenyi Biotec FeraTrack SPIO / 2 ml culture medium) was chosen according to cell viability and manufacturer’s instructions. SPIO uptake was confirmed using a Prussian blue iron assay kit (Millipore Sigma) according to manufacturer’s instructions. Day 30 SPIO-labeled cultures were encapsulated in neural ribbons plus Fluorescein-Dextran (500 kDa, 1:30 in alginate) and shipped on ice overnight as described above. Collagen phantom slabs were generated from 5 mg/ml PureCol EZ gel Type 1 Collagen solution (Advanced Biomatrix #5074) at RT by plating the solution in 3 cm dish. 60 μm diameter neural ribbons were manually severed to single 3 – 4 mm segments containing ~5,000 cells. Single ribbon segments were individually drawn into a 26G custom made curved needle Hamilton syringe with a ~45 degree bend, ~2 mm proximal to the bevel and injected by hand into collagen phantoms. Fluorescein-Dextran in neural ribbon segments was visualized using a Leica M165 FC stereo microscope.

### Rat cervical spinal cord hemi-contusion injury and transplants

For the rat SCI model, we induced cervical C4 hemi-contusion injuries as previously described (Mondello et al., 2015), wherein C4 contusion injuries were made using the electromagnetic spinal cord injury device (ESCID). Moderate lesions were made in 8-weeks-old female Long-Evans rats (N = 6 animals). In short, right-sided hemi-laminectomy was performed at C4 level in anesthetized animals that were then placed in a spinal frame to clamp the lateral processes of C3, C5. The ESCID electromagnetic probe was positioned at the surface of the dura with an initial sensing force followed by rapid displacement (0.8 mm with 14 ms dwell). Muscles were then sutured in layers and the skin was clipped. One week prior transplantation of human cells, the immunosuppressant tacrolimus was administrated at the dosage of 3.4 mg/kg/day through gradual release from subcutaneous implanted pellet (Innovative Research of America). Near the time of cell transplantation, 60 μm diameter neural ribbons containing day 30 human spinal neurons were shipped and received in Houston one day prior to injections. Constructs were suspended in fresh N2B27 and maintained in a humidified tissue culture incubator overnight. Cell number and viability was determined with an automatic counter (Invitrogen) before transplantation. For transplants, the spinal cord was re-exposed at injury level 15 days post-injury in anesthetized animals. To stabilize the animal, the lateral process of C3 was secured in a custom spinal frame. For delivery of high dose cell suspensions (N = 2 animals), alginate was dissolved using 1.6% sodium citrate and cells collected by centrifugation. 200,000 cells were delivered into the injury cavity by pulled glass pipette connected to Kopf micromanipulator in 2 μl of 5 mM Glucose in HBSS vehicle with the addition of 20 ng/ml FGF and EGF for neurotrophic support. Neural ribbon segments (N = 4 animals, ~5,000 cells per segment) were individually drawn into a 26G curved needle Hamilton syringe with a ~45 degree bend ~2 mm proximal to the bevel. The syringe was placed in a Kopf micromanipulator attached to the spinal frame and positioned 0.5 mm lateral to midline ipsilaterally. The syringe tip was used to puncture the dura, and advanced to a depth of 0.7 mm. The single ribbon segment was injected over 1 min in 5 μl of HBSS per animal. The needle was maintained in position for 2 min dwell, followed by 100 μm retraction and a second 2 min dwell, and then was slowly retracted while monitoring for efflux of fluid or the ribbon segment onto the surface of the dura. After injection, the dura was sealed using a fibrin sealant made by mixing a 1:1 ratio of fibrinogen solution (20 mg/ml) with a bovine thrombin solution (10 U/ml) (Millipore Sigma). Muscle and skin layers were closed, and animals received appropriate post-operative care. For visualization of graft placement by MRI, one additional uninjured control animal was injected with a neural ribbon segment containing SPIO-labeled cells at the C4 level. All rat experimental protocols were approved by Houston Methodist Research Institute IACUC and carried out in accordance with relevant ethical guidelines and regulations. This includes animal comfort, veterinary care, methods and reasons for euthanasia, and materials and hiPSC-derived neural injections.

### Ex vivo spinal column MRI

Spinal columns were imaged in a 9.4 T ultra-high field vertical MRI scanner (Bruker) using an adjusted multi-echo T2 rapid acquisition refocused echo sequence with acceleration factor of 4, 10 averages and single excitation cartesian k-space scan trajectory. Pixel bandwidth of 40,761 Hz/px, time-to-repetition of 1800 ms, and time-to-echo of 25, 37 and 45 ms were used. For *in silica* characterization of the neural ribbon transversal relaxation (T2) signal, samples were imaged in 200 μL plastic tubes containing physiologic saline solution. For *in situ* imaging, in-plane resolution of 25 μm and slice thickness of 1 mm was used for axial acquisitions with anterior-posterior phase encoding and FOV of 12.5 mm. 13 × 21 μm resolution and 3 mm slice thickness were used for coronal and sagittal acquisitions with rostral-caudal phase encoding directions and FOV of 50 mm. For quantitative analysis *in situ*, T2 relaxation maps were generated pixelwise by applying a single exponential fit to image intensity measured over four time-to-echo values. Transplanted neural ribbon constructs were visualized in the rodent spinal column using Horos Medical Image Viewer (Horos Project, 2020). Raw images acquired at time-to-echo = 25 ms and computed T2 maps were considered. An inverse log-scale colormap, typically applied to imaging of bone perfusion, was chosen for enhanced visualization of the ribbons.

### Tissue processing and immunohistochemistry

Eight days after neural ribbons were grafted, rats were sedated with isoflurane and transcardially perfused first with cold 0.1 M PBS with 10,000 U of heparin followed by 4% paraformaldehyde in 0.1 M PBS (pH 7.4). Spinal cords were harvested for storage in 4% paraformaldehyde at 4°C overnight. The spinal cords were passed through a sucrose buffer gradient (10%, 20%, 30%; 24 h per solution) at 4°C for cryoprotection. Cords were then aligned longitudinally in a Tissue Tek OCT block (Sakura, Nederland) using dry ice, and stored at −80°C. 40 μm thick spinal cord sagittal sections were prepared using a Cryostar NX50 cryomicrotome (ThermoFisher). Serial sections of the entire spinal cord were then mounted onto positively charged slides (Fisher Scientific). For immunohistochemistry, slides were first blocked with 10% goat serum in 0.2% Triton-PBS (T-PBS) for 1 h at room temperature followed by primary antibody incubation in 1% serum, 0.2% T-PBS overnight at 4°C. Primary antibodies used were mouse anti-STEM121 (1:500; Takara), chicken anti-GFAP (1:2,000; Abcam), and rabbit anti-Synapsin 1 (1:200; Abcam). After three additional wash steps with 1% serum T-PBS (1-2 min/wash), slides were incubated with secondary antibodies in PBS for 1 h. Secondary antibodies used were goat anti-mouse AF488 (1:1000; Invitrogen), goat anti-chicken Cy5 (1:1000; Invitrogen), goat anti-rabbit AF568, (1:1000; Invitrogen). Slides were mounted with DAPI-Permount (Invitrogen) and imaged with Leica DMi8 confocal microscope.

### Experimental design and statistical analyses

Raw data were compiled in Microsoft Excel (v16.16.16) and exported to GraphPad Prism (v8.3.0) for plotting and statistical analysis. Data are reported as (mean ± s.e.m.) and analyzed using unpaired two-tailed t-test unless otherwise specified. Cells were manually counted in ImageJ to quantify immunofluorescence data. One-way ANOVA test was used to compare patch clamp electrophysiology experimental conditions. ****p<0.0001, ***p<0.001, **p< 0.01, *p<0.05, n.s. not significant (α = 0.05). Power analysis was not performed for grafting studies. Detailed information for each experiment is provided in figure legends.

***Figures.*** Figures for this manuscript were made in Keynote (v9.2.1) and Canvas Draw (v4.0.1). Data plots were generated using GraphPad Prism (v8.3.0) or Microsoft Excel (v16.16.19).

## Results

### Generation of cervical SMNs from hiPSCs using trunk-biased developmental neuromesodermal progenitors

Spinal motor neurons (SMNs) were derived from the African American hiPSC line F3.5.2 (Chang et al., 2015; Tomov et al., 2016) in three stages (**Fig. 1** **and** **2**), here referred to as (1) neural induction, (2) caudo-ventral patterning, and (3) neuronal maturation. For neural induction we generated neuromesodermal progenitors (NMps) that are bipotent intermediates preceding spinal cord neuroectodermal development (Lippmann et al., 2015; Kumamaru et al., 2018). Synergistic Wnt/β-catenin and FGF-mediated signaling pathways were stimulated using the small molecule CHIR 99021 and recombinant human FGF2/FGF8 proteins, respectively, in the context of dual SMAD inhibition with LDN 193189 and SB 431542 (Chambers et al., 2009; Chambers et al., 2012) during the first five days of differentiation. The GSK-3β inhibitor DAPT was also included in the growth medium during this time. NMp transition to spinal cord NSC rosettes (scNSCs) was trigged by switching to cervical patterning medium containing 100 nM of the caudalizing morphogen, retinoic acid (RA), and 200 nM of the ventralizing morphogen, Hh-Ag1.5, a potent agonist of Sonic hedgehog (**Fig. 1*A***). Continued exposure to cervical patterning medium yielded robust motor neuron progenitor (MNP) cultures by day 12 that were immunopositive for Nestin, OLIG2, TUJ1, and NeuN (**Fig. 1*B*-1*D***).

**Figure 1.**
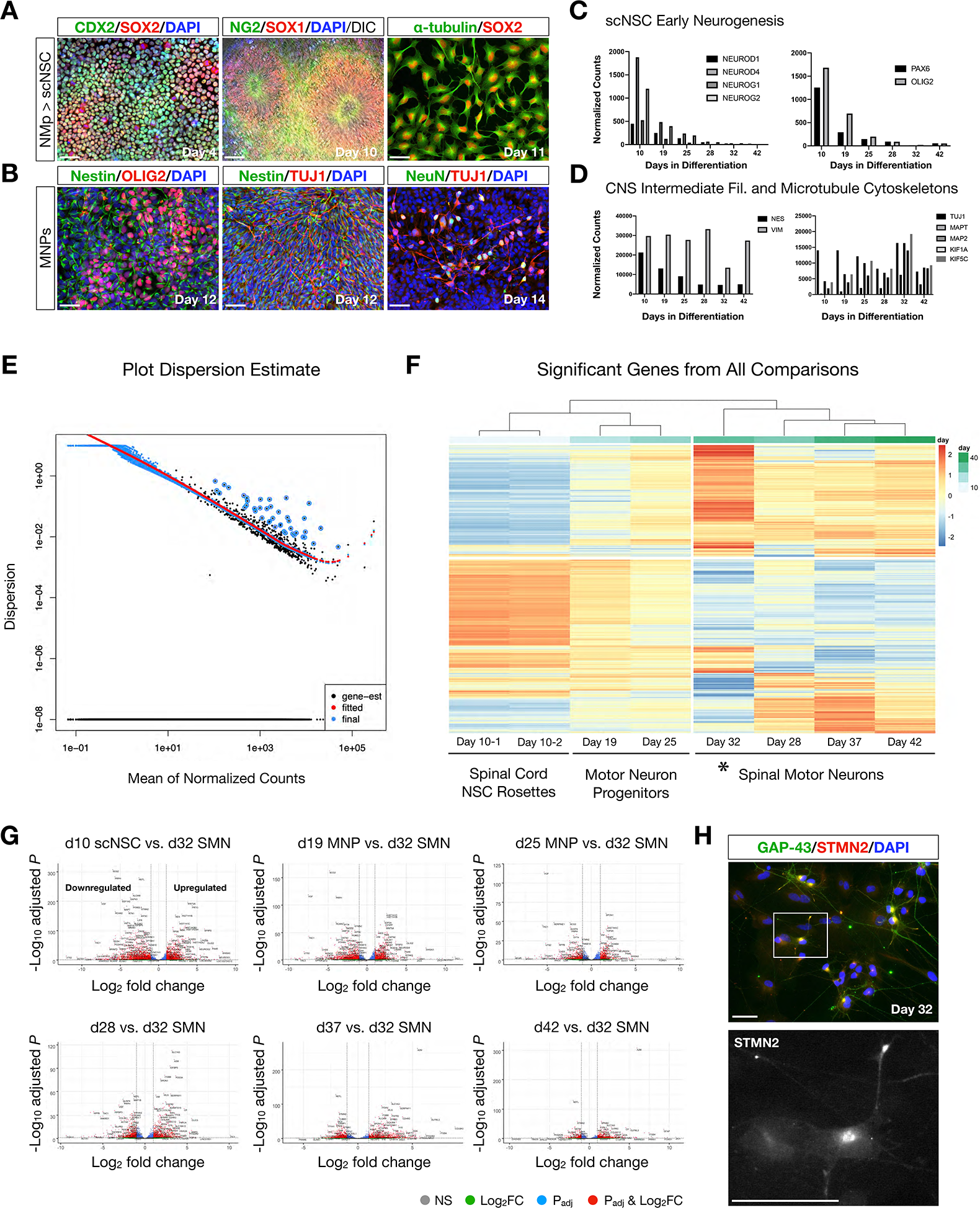
RNA-Seq bioinformatics analysis of scNSC neuronal differentiation. ***A***, CDX-2/SOX2 NMps generate NG2/SOX1 scNSC rosettes that can be reseeded as spinal cord neural progenitor cell adherent cultures (α-tubulin/SOX2). ***B***, Caudo-ventral patterning of scNSCs yields motor neuron progenitors (MNPs) expressing biomarkers Nestin, OLIG2, TUJ1 and NeuN. ***C***, Histogram of normalized counts by RNA-Seq reveals progressive downregulation of transcription factors involved in neurogenesis (left) and motor neuron-specific patterning (right) from scNSCs to a differentiated state. ***D***, Normalized counts of CNS-related intermediate filament (Nestin, Vimentin; left) and microtubule cytoskeleton proteins (right) by RNA-Seq. ***E***, Plot Dispersion Estimate of RNA-Seq samples. ***F***, Heatmap of significant genes from all comparisons for SMN differentiation from scNSC stage reveals four predominant clusters. Day 32 constitutes a gold standard for our SMNs. ***G***, Volcano plots of differentiation time points with reference to day 32 SMNs. ***H***, Top: IF of differentially expressed GAP-43 and STMN2 in day 32 adherent SMNs. Bottom: STMN2 characteristic perinuclear and growth cone localization (Klim et al., 2019). Scale bars are 50 μm.

**Figure 2.**
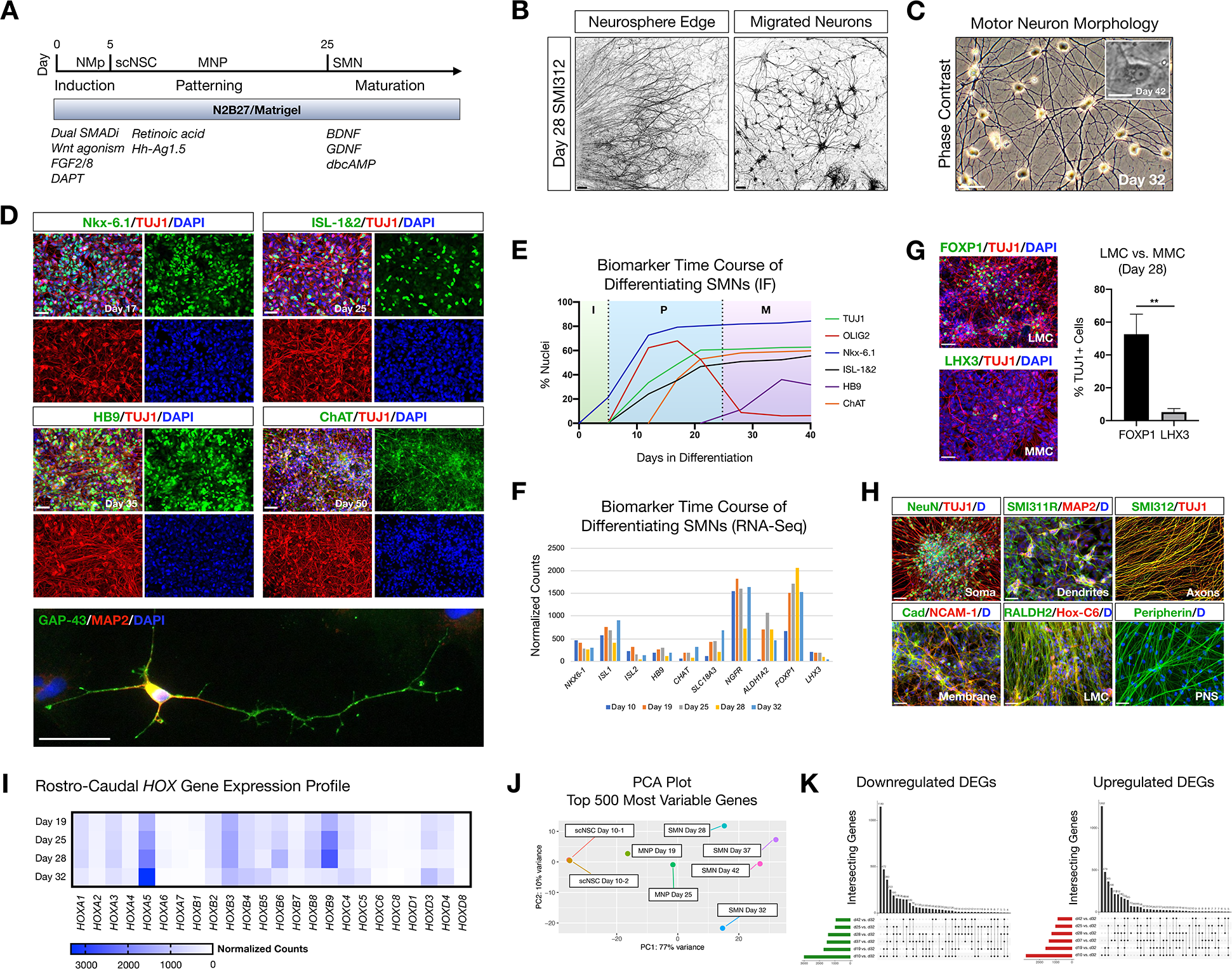
hiPSC-derived scNSCs generate SMNs with cervical spinal identity. ***A***, Overview of differentiation protocol used to generate cervically-patterned SMNs through NMp, scNSC and MNP stages. Small molecules and growth factors used in caudo-ventral patterning and maturation are listed. ***B***, Pan-axonal antibody SMI312 labeling of day 28 adherent neurosphere (NS) culture at NS edge (left) and more distal field of migrated neurons (right). SMI312 signal shown with inverted LUT. ***C***, Phase contrast images of day 32 SMN cultures. Inset is day 42 single SMN with characteristic trapezoidal shape and prominent nucleolus. ***D***, SMN hallmark biomarkers. Top: Motor progenitor spinal domain transcription factor Nkx-6.1 (day 17); SMN-related factors ISL-1&2 (day 25); Middle: SMN-specific HB9 (day 35); choline acetyltransferase enzyme ChAT (day 50); Cells are counterstained with TUJ1 and DAPI. Bottom: IF of single neuron. GAP-43/MAP2/DAPI (day 32). ***E***, Time course of biomarker expression in differentiating SMN cultures (I, induction; P, patterning; M, maturation). Percent of nuclei were counted from IF images. N = 3 fields were averaged per biomarker per timepoint (>100 cells per field). ***F***, Time course biomarker expression of differentiating SMN cultures by bulk RNA-Seq. ***G***, IF images and histogram of later motor column (LMC) SMN transcription factor FOXP1 (53 ± 12%; n = 349 cells) versus medial motor column (MMC) SMN transcription factor LHX3 (5 ± 2%; n = 304 cells). (two-tailed t test, t = 6.629, df = 4, **P = 0.0027). ***H***, Visualizing neuronal compartments and SMN regional specificity by biomarker. ***I***, RNA-Seq validation of cervical spinal cord identity by normalized *Hox* gene expression profile in differentiating SMN cultures. ***J***, PCA plot of 500 most variable genes during scNSC neuronal differentiation. ***K***, UpSet plots of intersecting downregulated (left) and upregulated (right) differentially expressed genes (DEGs) using individual time point comparisons in differentiation. Plot height represents the number of overlapping DEGs between the sets below marked with filled circles. Vertical lines connecting two or more dots denote the set of genes overlaps the connected sets. Data are reported as mean ± s.e.m. Scale bars are 50 μm, and 10 μm in (***C***) inset.

We used short-term (24 – 48 h) NS suspension cultures as a purification step prior to neuronal maturation and as a consistent technique for the statistical analysis of spinal motor neurogenesis. Plated NS cultures projected aligned axons radially outward along which neurons migrate to form distal 2-D adherent regions, visualized by the pan-axonal marker SMI312 (**Fig. 2*B***). At day 25, cultures were transitioned to and subsequently maintained in terminal differentiation medium containing 10 ng/ml BDNF, GDNF and 1 μM dbcAMP. Adherent neurons exhibited typical SMN morphology as assessed by trapezoidal soma shape, a prominent single nucleolus, and dendritic arborization (**Fig. 2*C***). The SMNs expressed a panel of characteristic biomarkers by IF time course (**Fig. 2*D*** **and** **2*E***). That is, the ventral motor neuron progenitor domain transcription factor NK6 homeobox 1 (Nkx-6.1), SMN-related ISL LIM homeobox 1 and 2 (ISL-1&2), SMN-specific homeobox 9 (HB9/*MNX1*), and choline acetyltransferase (ChAT). Growth associated protein 43 (GAP-43) localized to neuronal growth cone and branching structures (**Fig. 2*D***).

### Developmental hallmarks of NMp-derived cervical SMNs validated by RNA-seq and biomarkers

SMNs derived through NMp intermediates were validated by whole-transcriptome bulk RNA-Seq on a differentiation time course (**Fig. 1*E*-1*G*** **and** **Fig. 2*F***). SMNs primarily expressed lateral motor column (LMC) transcription factor FOXP1 (53 ± 12%, n = 349 cells) in comparison to the medial motor column (MMC) transcription factor LHX3 (5 ± 2%, n = 304 cells) that was downregulated over time (**Fig. 2*F*** **and** **2*G***) (Amoroso et al., 2013). We also verified general biomarker expression for neuronal compartments that are somatic (NeuN), dendritic (SMI311R/MAP2), axonal (SMI312) and membrane (Pan-Cadherin/NCAM-1), and phenotypic regional specificity to the cervical LMC (Hox-C6, RALDH2, Peripherin) by IF (**Fig. 2*H***). Cervical motor identity is supported by rostral-caudal collinear *Hox* gene expression profiles using bulk RNA-Seq (**Fig. 2*I***). PCA analysis plot of the top 500 most variable genes revealed the similarities between scNSC biological replicates, but a progressive dissimilarity between time points in differentiating cultures (**Fig. 2*J***). UpSet plots for downregulated (log_2_(fold change) < 0) and upregulated (log_2_(fold change) > 0) DEGs (P_adj_ < 0.05) enable visualization of multiple set intersections similarly to Venn or Euler diagrams (**Fig. 2*K***). However, unlike Venn or Euler diagrams, UpSet scales appropriately when comparing three or more groups and also maintains correct intersectional area proportions. Differentiating samples were compared to day 32 SMNs and the trend was the same for both downregulated and upregulated DEGs.

We interrogated maturation and functional hallmarks of the SMNs using numerous methodologies (**Fig. 3**). At day 35 in differentiation, SMNs were immunopositive for axon initial segment (AIS) proteins Ankyrin-G (AnkG) and βIV-Spectrin (**Fig. 3*A***). Immunostaining occurred as short segments along TUJ1 neuronal filaments distal to the soma for both AnkG (32 ± 2 μm) and βIV-Spectrin (34 ± 1 μm; mean ± s.e.m.) (**Fig. 3*B***). Time course RNA-Seq analysis of genes encoding the voltage-gated ion channels Nav1.6 (*SCN8A*) and Kv1.2 (*KCNA2*) additionally supports ongoing maturation in these cultures (**Fig. 3*C***). As well, robust mitochondrial transport was observed in neurites using live-cell imaging with MitoTracker Green FM.

**Figure 3.**
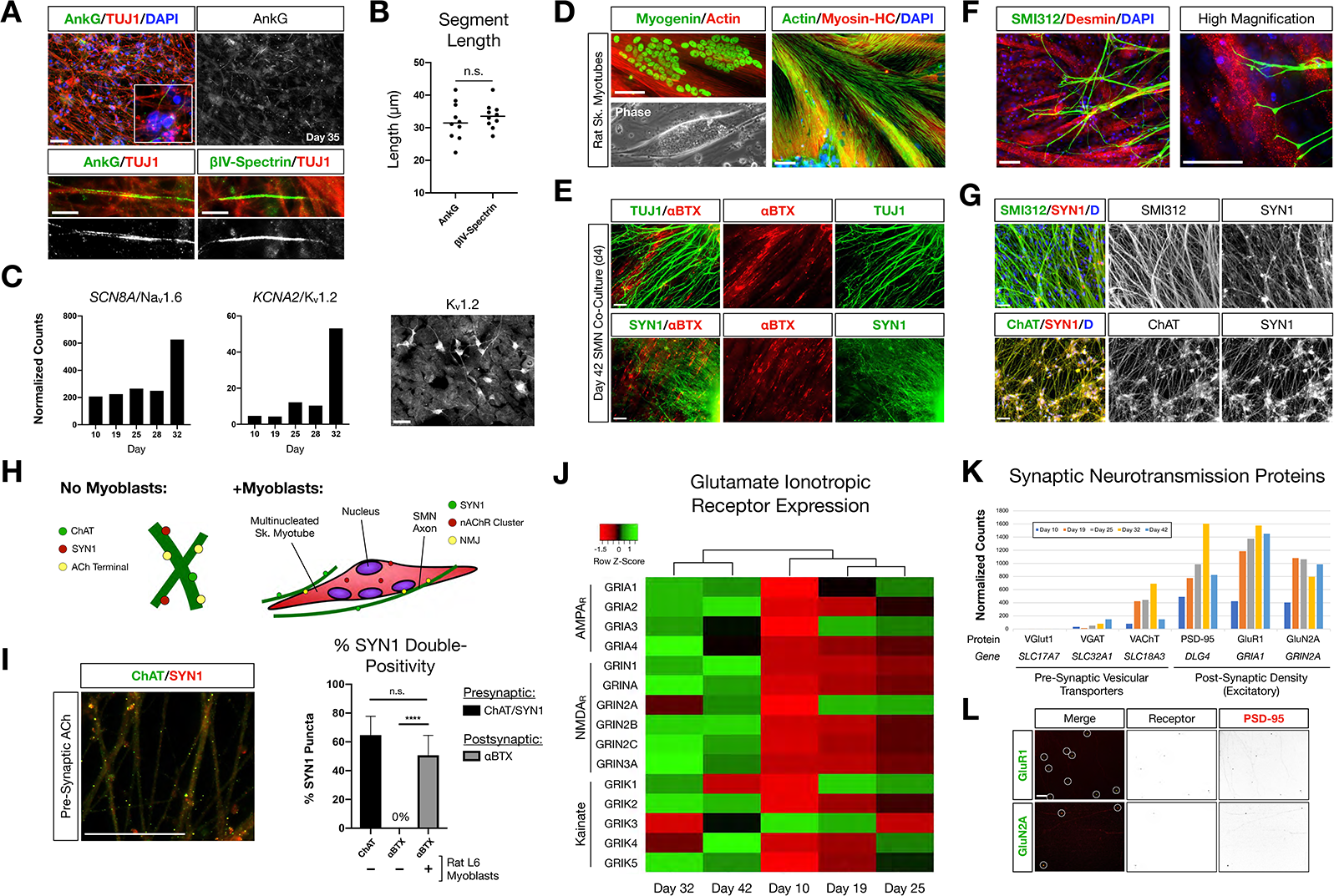
NMp-derived cervical SMNs form AIS, innervate myotubes and develop characteristic synaptic profiles. ***A***, Maturing SMN-NS cultures exhibit short segments immunopositive for AIS proteins Ankyrin-G (AnkG) and βIV-Spectrin at day 35. Top: AnkG/TUJ1 counterstained with DAPI (nuclei). High-magnification image depicts AnkG+ staining distal to the soma. Bottom-left: AnkG immunopositive region along TUJ1+ filaments. Bottom-right: βIV-Spectrin immunopositive region along TUJ1+ filaments. ***B***, Length of AnkG (32 ± 2 μm) and βIV-Spectrin (34 ± 1 μm) segments. (two-tailed t test, t = 0.7082, df = 18, P = 0.4879, n = 10 segments measured per antibody). ***C***, Time course normalized expression of AIS-related voltage gated ion channels Nav1.6 and Kv1.2 by RNA-Seq. Kv1.2 IF image on right. ***D***, Rat L6 multinucleated skeletal myotubes for NMJ formation assays. ***E***, Day 4 co-culture of SMN-NS (day 42) seeded onto rat myotubes. Top: TUJ1+ filament termination onto αBTX+ junctions (nAChR). Bottom: SYN1/αBTX. ***F***, SMI312 axon termination in Desmin+ myotubes resembling early motor end plates. ***G***, Co-localization of cholinergic terminals (ChAT, bottom) with SYN1 puncta along axons (SMI312, top). ***H***, Cartoon diagram of ChAT/SYN1 pre-synaptic ACh terminals and nAChR post-synaptic machinery. ***I***, IF image and histogram of ChAT/SYN1 (64.7±13.1%, n = 118 puncta) and SYN1/αBTX co-localization in the absence (0%, n = 48 puncta) or presence (50.7 ± 13.8%, n = 69 puncta) of L6 myotubes. (two-tailed t test, t = 10.36, df = 14, ****P < 0.0001). N = 8 fields were averaged for each condition. ***J***, Heatmap of glutamate ionotropic receptor expression (AMPAR, NMDAR, Kainate) during SMN differentiation. ***K***, Histogram of normalized gene expression for pre-synaptic vesicular transporters (VGlut1 excitatory, VGAT inhibitory, VAChT cholinergic) and post-synaptic density proteins/excitatory receptors. ***L***, Co-localization of AMPAR GluR1 (top) and NMDAR GluN2A (bottom) with scaffolding protein PSD-95. Individual channels provided as inverted LUT. Data are reported as mean ± s.e.m. Scale bars are 50 μm, and 10 μm in (***A***) high magnification images (bottom).

Functional ability to form neuromuscular junctions (NMJs) was assessed between species by generating multinucleated skeletal myotubes from rat L6 myoblasts (**Fig. 3*D***). Co-culture of SMN-derived NS (day 38) with rat myotubes for four days resulted in marked co-localization of TUJ1 and Synapsin 1 (SYN1) neuronal filaments with α-bungarotoxin (αBTX), a high-affinity antagonist of nicotinic acetylcholine receptors (nAChR) (**Fig. 3*E***). SMN axons labeled with SMI312 terminated abruptly onto myotubes and formed structures resembling early motor end plates (**Fig. 3*F***). We observed co-localization of ChAT with SYN1 puncta along SMN axons as cholinergic pre-synaptic terminals (64.7 ± 13.1%, n = 118 SYN1 puncta) (**Fig. 3*G*-3*I***). Post-synaptic αBTX did not co-localize with SYN1 puncta in SMN mono-cultures (**Figs. 3*H*** **and** **3*I***). SMN activity in the spinal cord is influenced by descending cranial motoneuron (MN) fibers via glutamatergic neurotransmission. We therefore investigated the expression of glutamate ionotropic receptors by the SMNs (**Fig. 3*J*-3*L***). Clustering analysis revealed that SMNs cluster separately from scNSCs and MNPs with progressive activation of AMPA, NMDA and kainate receptors from scNSCs through to SMNs as a general trend (**Fig. 3*J***). In accordance with ChAT and SYN1 pre-synaptic co-localization, SMNs primarily expressed the acetylcholine vesicular transporter VAChT versus inhibitory (VGAT) or excitatory (VGlut1) vesicular transporters. However, the excitatory receptor post-synaptic density scaffolding protein PSD-95 (*DLG4*) was highly expressed as with the AMPA receptor subunit GluR1 (*GRIA1*) and the NMDA receptor subunit GluN2A (*GRIN2A*) (**Fig. 3*K***). GluR1 and GluN2A were trafficked to the membrane and co-localized with PSD-95 (**Fig. 3*L***).

### Characterization and co-encapsulation of sub-populations of spinal interneurons and glia present in scNSC differentiated SMNs

Ventral and dorsal spinal INs as well as astrocytes are indispensable for synaptogenesis support SMN network formation and connectivity. The differentiating SMN cultures analyzed by RNA-Seq and IF generated broad ventral as opposed to dorsal identity as analyzed by mRNA normalized counts of characteristic progenitor domain and neuronal subtype transcription factors (**Fig. 4*A***) (Sagner and Briscoe, 2019). By IF, we quantified the proportions of nine generic classes of spinal neurons in day 58 cultures that are Nkx-2.2 (V3, ~2%), FOXP1+ISL1 (MN, ~52%), CHX10 (V2a, ~12%), GATA-3 (V2b, < 1%), PAX2 (V0/V1 or dI4/dI6, ~8%), LBX1 (dI4-dI6, 0%), TLX3 (dI3, ~1%), LHX9 (dI1, 0%), and BRN3A (sensory neurons from neural crest or dorsal INs, ~25%), and provide representative images for CHX10, PAX2 and BRN3A (**Fig. 4*B*** **and** **4*C***). Low levels of LBX1, TLX3 and LHX9 in the presence of Peripherin indicate that BRN3A+ cells are neural crest-derived sensory neurons that can be patterned through NMps. We also identified genes for oligodendroglial and astroglial lineages in differentiating SMN cultures (**Fig. 4*D*** **and** **4*E***). GFAP+ astrocytes were not observed until day 40 in differentiation, consistent with late waves of gliogenesis subsequent to an initial neurogenesis phase in human gestation. In adherent cultures, IN subtypes developed in close proximity as small clusters, and this configuration was retained within neural ribbon-encapsulated constructs (**Fig. 4*F***). Cultures encapsulated prior to day 40 do not contain GFAP+ astrocytes since these cells are not yet present.

**Figure 4.**
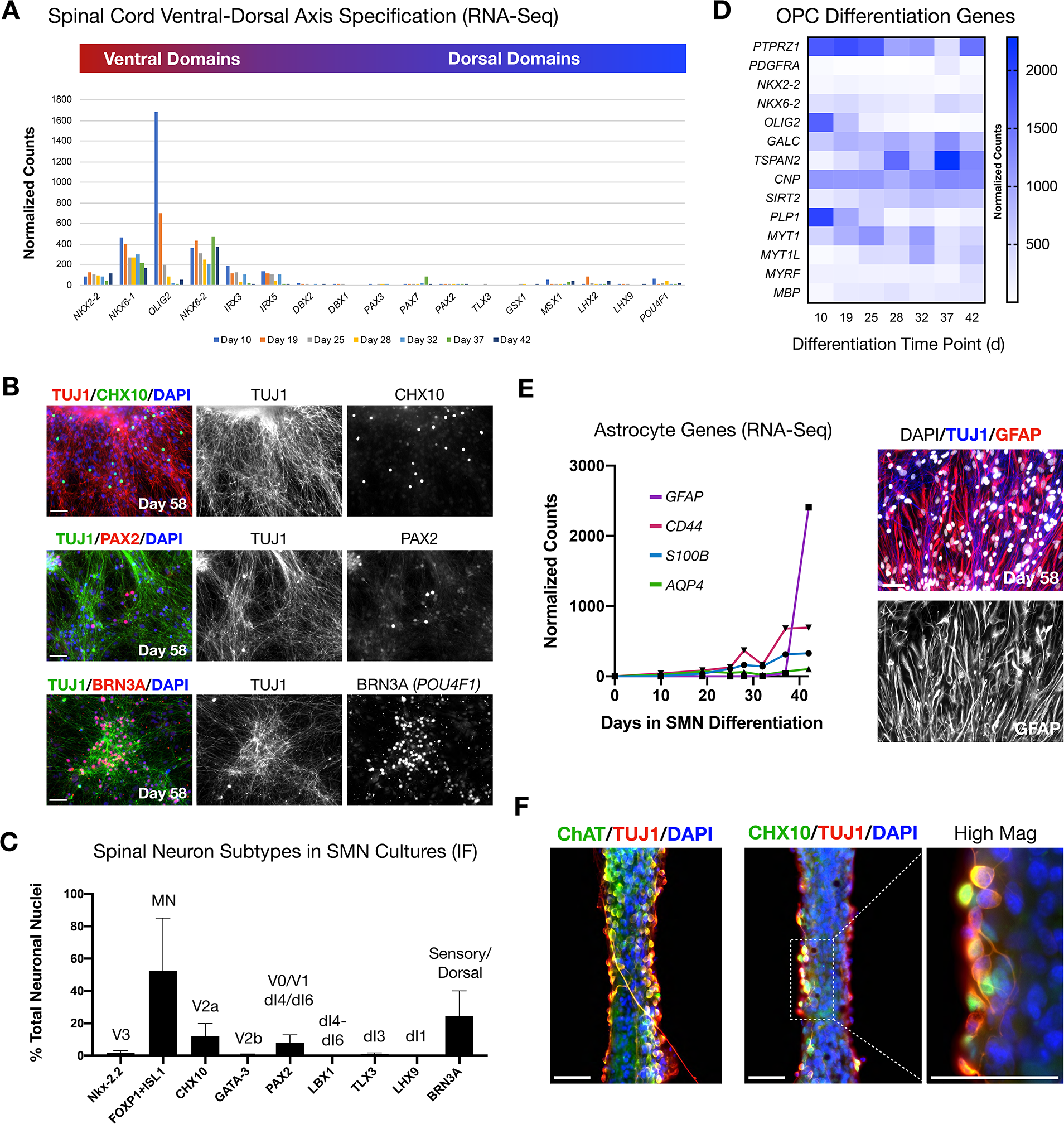
Quantifying sub-populations of interneurons and glia that emerge in SMN cultures. ***A***, Histogram of RNA-Seq normalized counts for ventral (red) and dorsal (blue) spinal cord domain transcription factors over time in differentiating SMN cultures at day 58 (Sagner and Briscoe, 2019). ***B*** Representative IF of IN subtypes in differentiating SMN cultures, quantified in (***C***). ***C***, IF quantification of spinal INs subtypes (% of total neuronal nuclei counted; mean ± s.e.m.). Domains shown: Nkx-2.2 (V3), FOXP1+ISL1 (MN), CHX10 (V2a), GATA-3 (V2b), PAX2 (V0/V1 or dI4/dI6), LBX1 (dI4-dI6), TLX3 (dI3), LHX9 (dI1), BRN3A (neural crest-derived sensory, or dorsal IN). ***D***, Heatmap of differentiation genes for OPCs that originate from the motor neuron progenitor ventral domain in differentiating SMN cultures. ***E***, RNA-Seq normalized counts over time for astrocyte biomarkers *GFAP*, *CD44, S100B* and *AQP4* in differentiating SMN cultures (left). A wave of gliogenesis occurs at late stage timepoints; GFAP/TUJ1 mixed culture (day 58). Cells are counterstained with DAPI (right). ***F***, ChAT (left) vs. CHX10 (right) immunostain in neural ribbons counterstained with TUJ1. High magnification image of CHX10 cluster (rightmost). Scale bars are 50 μm.

### NMp-derived cervical spinal neurons develop regular spiking patterns and glutamate-responsive bursting

Neuronal firing was examined by calcium imaging as a proxy measure, and by MEA extracellular recordings (**Fig. 5**). For adherent cultures, we used the cell-permeant calcium-sensitive dye, Fluo-4 AM and measured calcium transients in the soma (**Fig. 5*A***, day 32) and in three regions of interest (ROIs) along the neurites (**Fig. 5*B***) demonstrating spontaneous activity. For MEA analysis we seeded day 28 neurospheres (NS) onto single 60-electrode arrays (12-15 NS/array) in terminal differentiation medium (**Fig. 5*C***, day 34) and maintained adherent cultures to day 52. The firing activity recorded using MEAs is described for day 35 and day 52 time points (**Fig. 5*D*-5*K***). Day 35 cultures exhibited high and irregular spontaneous firing (**Fig. 5*D*** **and** **5*H***). By contrast, day 52 cultures displayed lower and more regular spontaneous firing rates (**Fig. 5*E*** **and** **5*H***; mean firing rate, MFR) (**Fig. 5*K*** **and** **5*I***). Notably, in both cultures, addition of 50 μM glutamate to the extracellular medium increased neuronal firing and produced robust bursting activity (day 35 ***P = 0.0007, day 52 **P = 0.0048). Glutamate-stimulated bursting was more pronounced and became qualitatively rhythmic in day 52 SMNs and this effect persisted for the entire duration of the experiment (**Fig. 5*K***).

**Figure 5.**
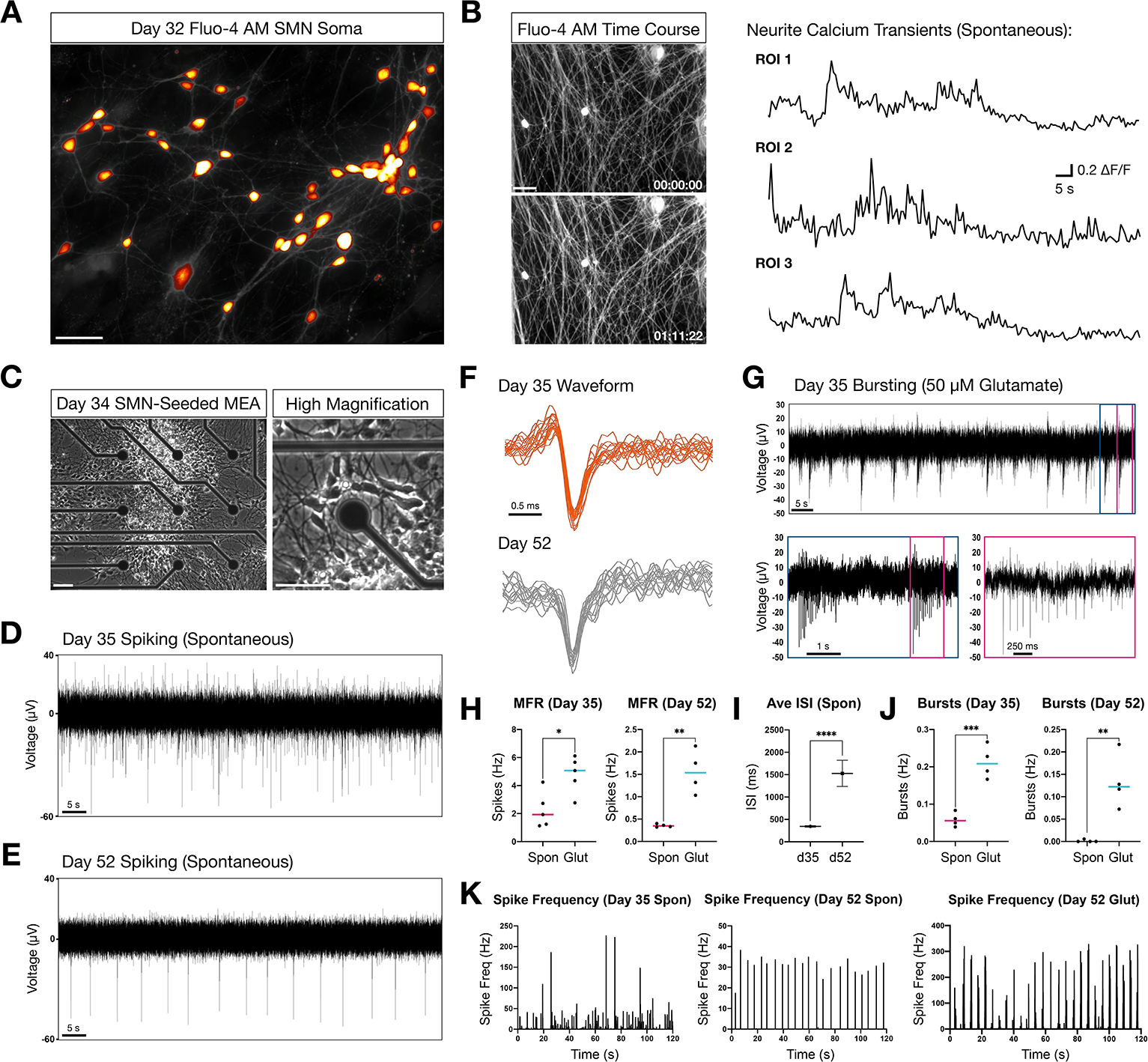
Firing properties of NMp-derived adherent spinal neurons by calcium imaging and microelectrode arrays. ***A***, Calcium dye Fluo-4 AM somatic signal in adherent SMNs (day 32). ImageJ Smart LUT. ***B***, Time course Fluo-4 AM fluorescence panels and ΔF/F quantification from three ROI in neurites. ***C***, Phase contrast images of SMN-seeded MEAs at day 34 in differentiation (seeded day 28). ***D-E***, Representative traces from single MEA channel depicting spontaneous SMN voltage spikes recorded at day 35 (***D***) and day 52 (***E***) in differentiation. *F*, Representative overlay of spontaneous spike waveforms at day 35 and day 52 from traces in (***D*, *E***). ***G***, Representative single electrode bursting in day 35 cultures stimulated with 50 μM glutamate. Three time scales are shown. Pink box depicts spikes within a single burst. ***H***, Quantification of mean firing rate (MFR, Hz) in day 35 and day 52 SMNs. Spontaneous firing is compared to 50 μM glutamate stimulation. (two-tailed t test; day 32 t = 3.096, df = 8, *P = 0.0148; day 52 t = 4.968, df = 6, **P = 0.0025). ***I***, Average inter-spike interval (ISI) in day 35 versus day 52 SMNs under spontaneous conditions. (t = 10.52, df = 499, ****P < 0.0001). *J*, Single electrode burst rate (Hz) in day 35 versus day 52 SMNs. (day 32 t = 6.426, df = 6, ***P = 0.0007; day 52 t = 4.356, df = 6, **P = 0.0048). ***K***, Spike frequency (Hz) versus time (s) in day 35 versus day 52 SMNs. Also shown is spike frequency for day 52 SMNs stimulated with 50 μM glutamate is shown (rightmost plot). Scale bars are 50 μm.

### Spinal neuron process alignment can be directed within hydrogel neural ribbons

We applied a hydrogel-based platform referred to as neural ribbons (Olmsted et al, 2020) to reproducibly encapsulate MNPs and SMNs, and optimize methodologies to direct and image cytoarchitecture configurations (**Fig. 6*A***). Hydrogel composition is dependent on alginate biopolymer crosslinking and includes addition of Type 1 collagen as exogenous ECM to establish interpenetrating networks (IPNs). The rapid extrusion and concomitant encapsulation of cells within 3D geometries was facilitated with 100 mM CaCl_2_ (**Fig. 6*B***). Cell-free and cell-laden ribbons each retained the circular cross-section and diameter of the needle tip template (60, 100, or 150 μm inner diameter) in CaCl_2_, but swelled by ~33% in culture medium. Sodium alginate (1.5% in NaCl) was further functionalized with bioactive peptide RGD that is used to promote cell adhesion and cell-cell interactions within neural ribbons (RGD-Alginate + Collagen, or RGD-Col). For immobilization, long-term culture and imaging we embedded neural ribbons in fast-gelling collagen solution in the viewing area of a glass bottom dish. Neurite extension within 60 μm neural ribbons was constrained along the ribbon longitudinal axis by addition of 4 μg/ml Aggrecan (chondroitin sulfate proteoglycan, CSPG) within the embedding medium. CSPG is inhibitory to axon penetration. Day 20 MNPs were encapsulated as small triturated NSs (**Fig. 6*C***) or larger intact aggregates (**Fig. 6*D***) at 1 × 10^8^ cells/ml concentration equivalent and cultured for up to 12 days. Cultures were transitioned to neuronal maturation at day 25 by changing from cervical patterning to terminal differentiation medium according to our protocol (**Fig. 2*A***). We achieved neurite anisotropic longitudinal alignment within neural ribbons by triturated NS encapsulation and collagen/Aggrecan embedding (**Fig. 6*C***) and observed expression of lateral motor column SMN phenotypic biomarkers, FOXP1 and ChAT, that co-localized with TUJ1 (**Fig. 6*D***). To investigate the physiological relevance of neural ribbon MNP encapsulation and SMN differentiation, we probed for pre- and post-synaptic proteins SYN1 and PSD-95, respectively, and observed robust co-localization in neural ribbons as puncta by day 10 post-encapsulation (**Fig. 6*E***). Neural cell borders were visualized by co-staining with NCAM-1. By excluding Aggrecan CSPGs from the embedding medium, unconstrained neurites projected laterally outward from the neural ribbon body in addition to internal growth, visualized in live cultures using MitoTracker Green FM (day 10 after encapsulation; **Fig. 6*F***). Neurite outgrowth further enabled contact between adjacent neural ribbons (**Fig. 6*G***).

**Figure 6.**
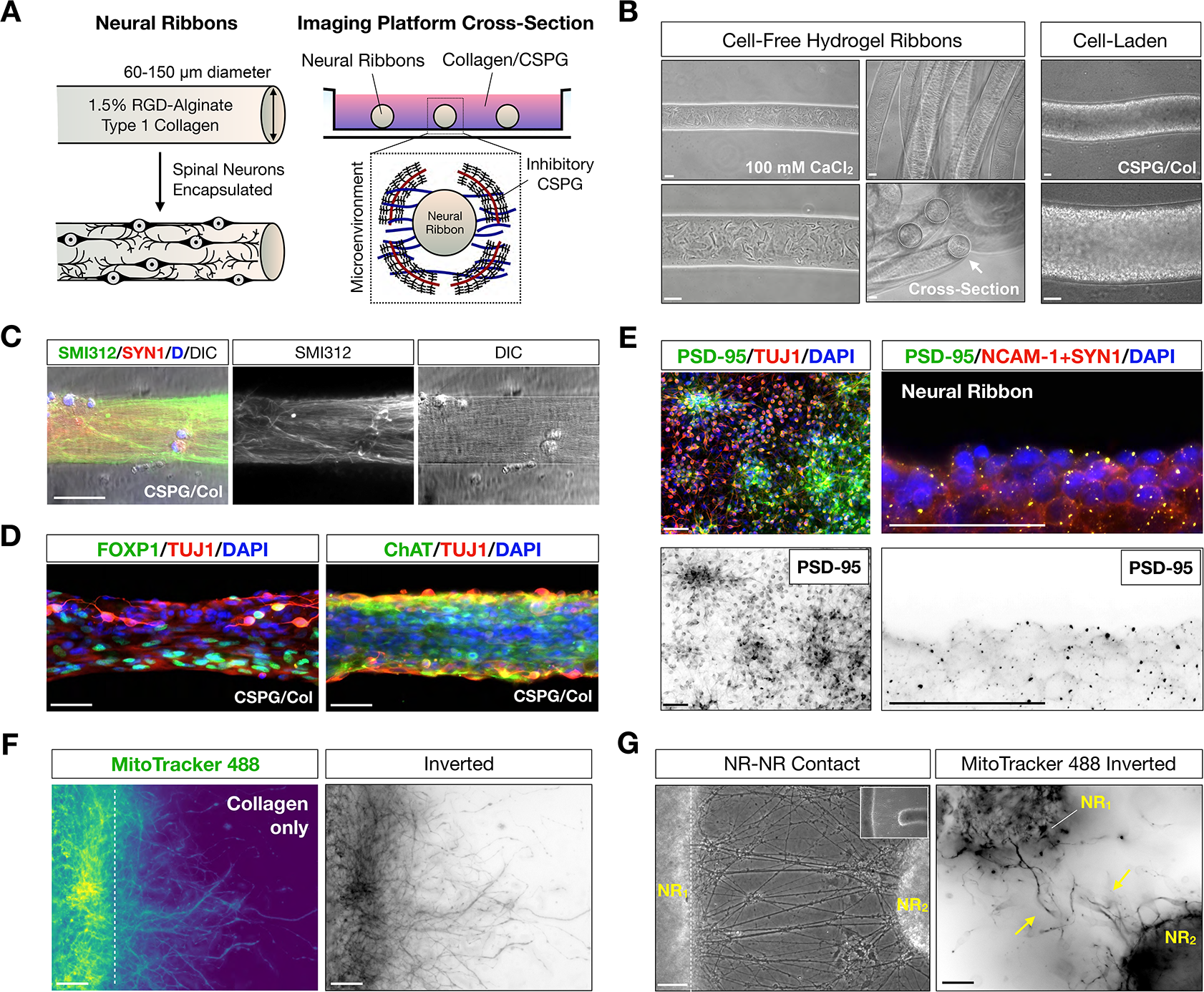
Directing spinal neuron configurations encapsulated in neural ribbons. ***A***, Cartoon diagram of SMN encapsulation in hydrogel neural ribbons and ECM embedding platform for imaging studies (glass bottom dish). Inhibitory CSPG components in collagen embedding matrix constrain axodendritic growth unidirectionally. ***B***, Phase contrast images of 1.5% RGD-functionalized alginate-based ribbons formed by extrusion into 100 mM CaCl_2_ crosslinking solution (left). Long ribbons adopt concentric configurations (middle-top) and have circular cross-sections (middle-bottom). Extrusion-based technique enables high-density encapsulation of neural cells in RGD-Col hydrogel ribbons. ***C***, Directed anisotropic SMN axonal configurations templated in neural ribbon platform. Ribbon stained with SMI312 (green), SYN1 (red), DAPI (blue). DIC image is shown. ***D***, SMN phenotyping within neural ribbons reveals retained expression of FOXP1 (left) and ChAT (right). Ribbons counterstained with TUJ1 and DAPI. ***E***, Glutamate receptor scaffolding protein PSD-95 expression in adherent SMN cultures (left) and co-localization with SYN1 in neural ribbons (right) at day 10 post-encapsulation. Additional NCAM-1 stain highlights cell boundaries. ***F***, Exclusion of inhibitory CSPG in collagen embedding matrix permits neurite outgrowth laterally from the neural ribbon body. Neural ribbon is stained with MitoTracker 488 (live cell) shown in green (left) and inverted LUT (right). ***G***, Neurite outgrowth enables connectivity between adjacent neural ribbons. Scale bars are 50 μm.

### Spinal neurons recovered from neural ribbons retain action potential firing

We analyzed SMNs treated with BDNF and GDNF neurotrophic factors using slow-release PLGA microbeads to test their effect on neuronal maturation (**Fig. 7*A***). At day 23, MNPs were passaged as aggregates and encapsulated in 60 μm neural ribbons (ProNova alginate + collagen, or NovaCol) lacking RGD peptide, or were used to generate MNP-NS cultures without encapsulation. MNP neural ribbons and MNP-NS were cultured in suspension for 48 h. On day 25, MNP aggregates were recovered from neural ribbons by dissolving alginate in 1.6% sodium citrate and seeded onto Matrigel-coated cover glass (**Fig. 7*B*** **and** **7*C***) in parallel with non-encapsulated MNP-NS. Cultures were further subject to SMN terminal differentiation and maintained up to day 52. To compare the effect of neural ribbon encapsulation on NMp-derived cervical SMNs on passive membrane properties and action potential firing at the single cell level, we performed whole-cell current clamp electrophysiology recordings (**Fig. 7*D*-7*F***). The resting membrane potential, input resistance and cell capacitance did not differ between 32-35 and 45-52 day SMNs (**Fig. 7*E***, top). We used direct current injections to hold SMNs at a potential of −65 mV, and then applied 5 ms long current steps to measure the rheobase, the minimum current amplitude that evokes an action potential, and the action potential threshold and peak (**Fig. 7*E***, bottom). Longer current injections (500 ms) were used to monitor the firing rate of the recorded cells in response to supra-threshold current injections. The data showed that all these parameters were similar in 32-35 and 45-52 day SMNs, suggesting that the excitability of these cells does not change substantially over this time interval. Consistent with these findings, we did not detect any significant difference in the frequency/current (f/I) relationship at these two time points (**Fig. 7*F***, left). The passive membrane properties and cell excitability properties of SMNs at these two different time points did not change in cultures treated with BDNF and GDNF (**Fig. 7*F***, middle) or in neural ribbon cultures (**Fig. 7*F***, right). As a consequence, their f/I plots were also unchanged. These results are important because they indicate that neurotrophic factors and neural ribbons do not disrupt or alter the firing properties of individual spinal neurons.

**Figure 7.**
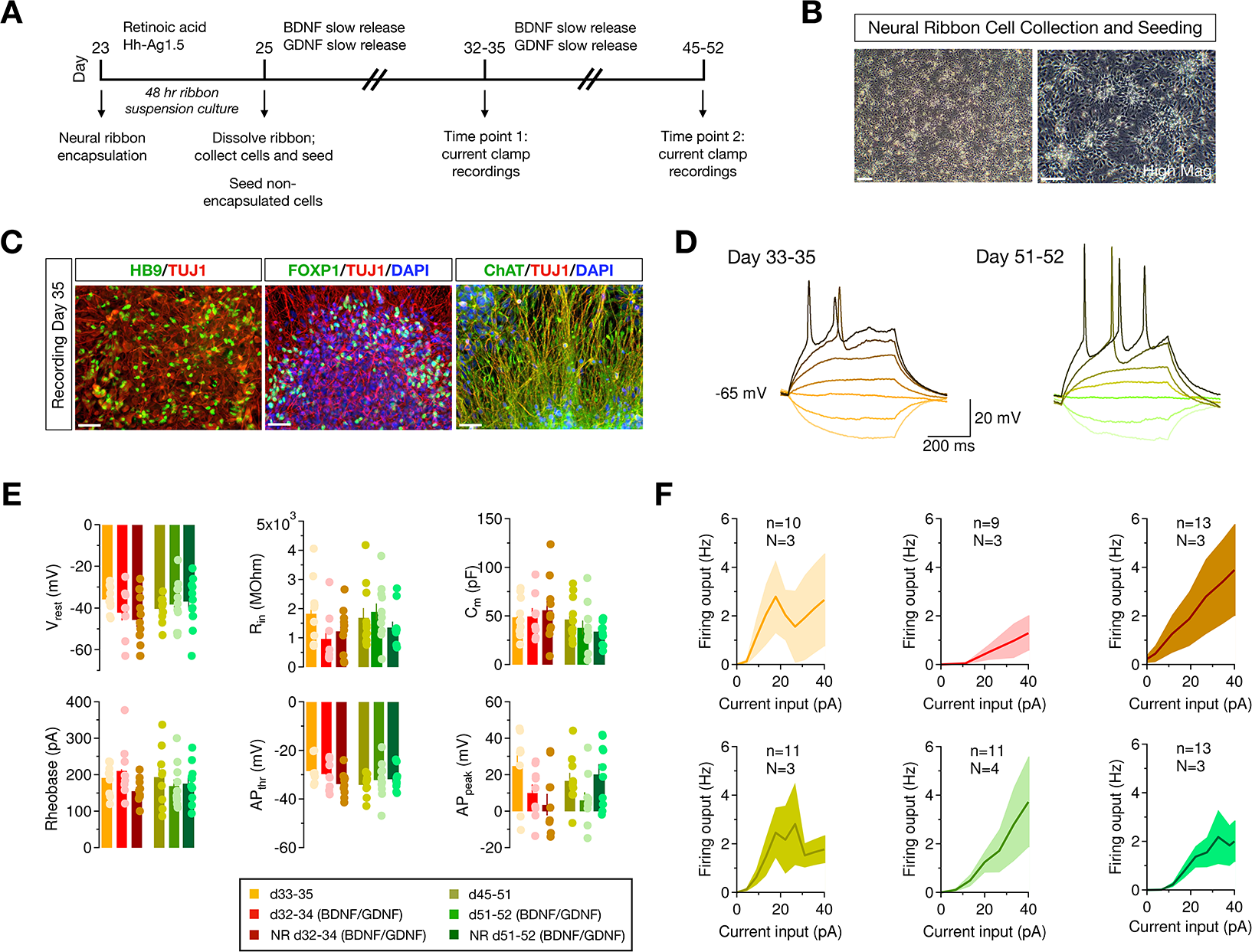
Neural ribbons preserve action potential firing properties. ***A***, Overview of the timeline used to perform the patch-clamp experiments. Differentiating SMN cultures were either encapsulated into neural ribbons (NR) for 48 h, collected, and seeded onto coverslips at day 25, or 48 h SMN-NS were seeded onto coverslips without NR encapsulation. Also tested was exposure to slow release of neurotrophic factors BDNF and GDNF from PLGA microbeads for neuronal maturation beginning at day 25. Current clamp recordings were performed at two time points (days 32-35 and days 45-52) across N = 3 - 4 differentiating cultures per condition (‘n’ denotes total number of cells recorded, whereas ‘N’ represents the number of culture batches used for these experiments). ***B***, Adherent SMN cultures 7 d after collection and seeding from NR culture. ***C***, SMN hallmark biomarkers at the day of recording (day 35) by IF. Scale bars for panels (***C*, *D***) are 50 μm. ***D***, Example traces of voltage responses to prolonged current step injections for time point 1 (left) and 2 (right). ***E***, Passive membrane properties (top: resting membrane potential, V_rest_, input resistance, R_in_, and membrane capacitance, C_m_) and active membrane properties (bottom: rheobase, action potential threshold AP_thr_, and action potential peak AP_peak_) for the six conditions tested. Dots represent the raw data collected from each cell (**Table 2**). No statistically significant difference was detected across samples by ANOVA. Sample key is provided (bottom). ***F***, Firing output (Hz) vs. current input (pA) for six conditions tested. Data represent mean ± s.e.m.

### Spinal neurons in 3D neural ribbons are glutamate-responsive interconnected networks

We employed an MEA system in conjunction with calcium imaging to track network formation in the neural ribbons (**Fig. 8**). SMNs were encapsulated in RGD-Col neural ribbons at day 28 in differentiation and maintained in terminal differentiation medium. At day 35, we performed non-adherent, acute recordings in three separate neural ribbons positioned over hexagonal arrays of MEA electrodes and immobilized under a coverslip. Recordings were acquired before and after addition of 50 μM glutamate. In one neural ribbon with the highest degree of firing, spiking activity was observed down the neural ribbon length and was significantly increased by glutamate stimulation in four of six electrodes (**Fig. 8*A*** **and** **8*B***). Spike raster plots in the two most active electrodes (C1, D4) are provided (**Fig. 8*C***), indicating the time point of glutamate addition. To assess the degree of connectivity with the neural ribbons, we quantified both intra-electrode bursting activity in addition to network burst (NB) events (**Fig. 8*D***). Burst rate was again increased by glutamate versus spontaneous conditions. We quantified total NB events and the percent of spikes contained within these events, that ranged from ~50 to 70%, where average NB duration was 1,011 ms. As additional validation of SMN spontaneous firing and connectivity in neural ribbons, we performed live-cell calcium imaging with Fluo-4 AM on encapsulated neurons (60 μm RGD-Col), in which linear arrangements of cells were visible (**Fig. 8*E***). Aggregates that retain neural ribbon shape following alginate dissolution in 1.6% sodium citrate (**Fig. 8*F***) exhibited retained rhythmic oscillations of somatic calcium transients (**Fig. 8*G***) and observable network bursting activity.

**Figure 8.**
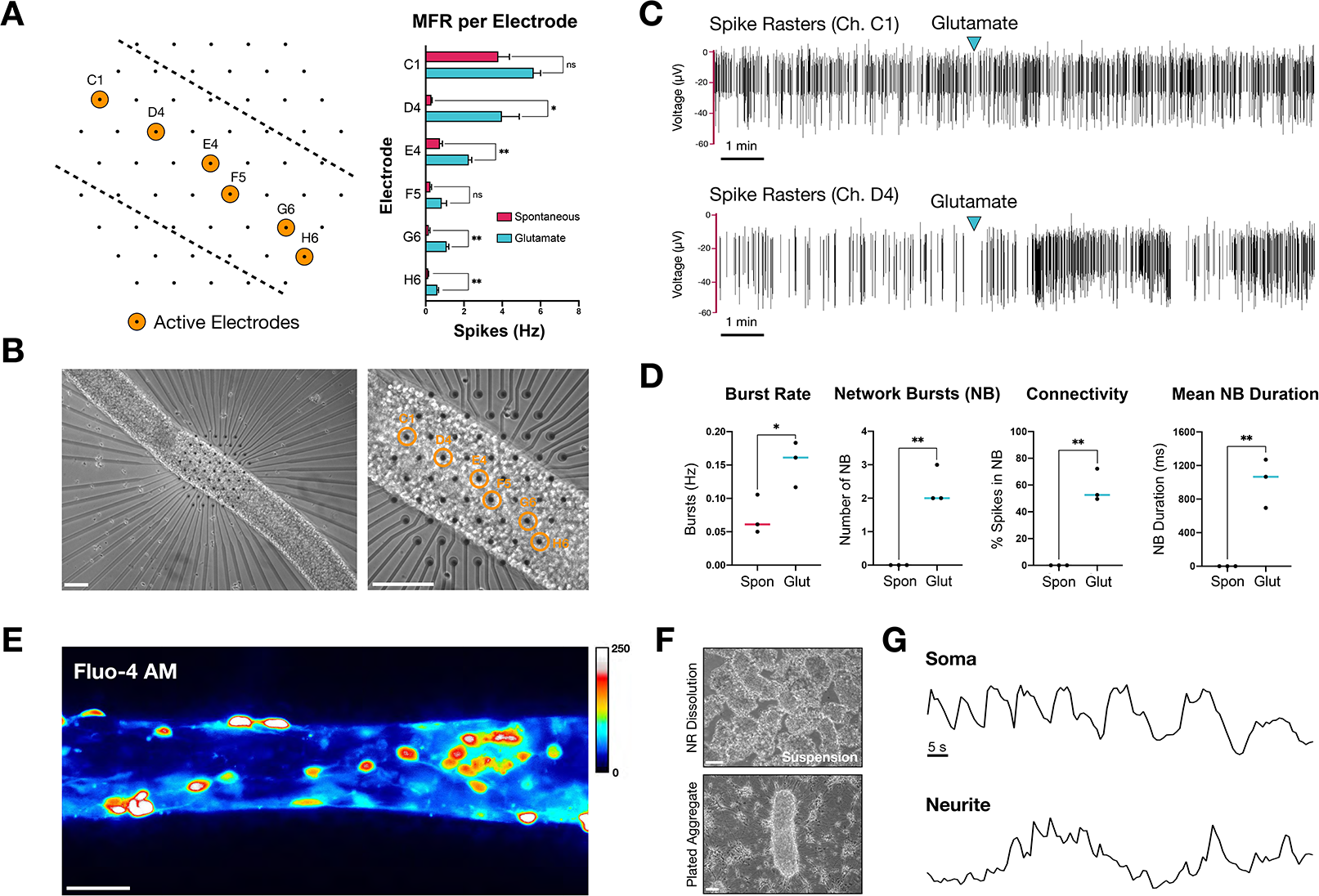
Glutamate-responsive functional activity in neural ribbons. ***A***, Left: schematic of neural ribbon recordings on hexagonal MEA array. Orange circles represent active electrodes. Right: histogram is mean firing rate (MFR, Hz) recorded in cell culture medium or medium supplemented with 50 μM glutamate. ***B***, Phase contrast images (day 35) corresponding to (***A***). ***C***, Spike rasters of electrodes with highest firing rate before and after addition of 50 μM glutamate. ***D***, Quantification (N = 3 neural ribbons) of burst rate (Hz; t = 3.143, df = 4, *P = 0.0348), number of network bursts per recording (NB; t = 7.000, df = 4, **P = 0.0022), connectivity (% spikes in NB; t = 8.212, df = 4, **P = 0.0012), and mean NB duration under spontaneous conditions versus 50 μM glutamate stimulation (t = 6.012, df = 4, **P = 0.0039). ***E***, Fluo-4 AM signal in 60 μm SMN neural ribbon (ImageJ Royal LUT). ***F***, Suspended neural aggregates recovered from 1.6% sodium citrate dissolution of alginate (top) and plating on fresh substrate (bottom, day 2). ***G***, Temporal oscillation of calcium transients in soma and neurites within plated aggregate from (***F***) plotted as ΔF/F. Scale bars 100 μm in (***B***), 50 μm in (***E*,*F***).

### Customized encapsulation of SMNs and oligodendroglia as purified co-cultures

We extended our neural ribbon technology to co-encapsulate SMNs with hiPSC-derived oligodendrocyte progenitor cells (OPCs) as pre-connected networks (**Fig. 9**). OPCs were differentiated similarly to SMNs up to day 20 but excluding DAPT during neural induction (**Fig. 9*A***). OPC differentiation proceeded through scNSC neuroectoderm (PAX6) and the motor progenitor spinal domain (Nestin/OLIG2), but were specified to oligodendrogenic progenitors by exposure to growth factors IGF1, FGF2, and PDGF-AA as well as thyroid hormone triiodothyronine (T3) at day 20 (Khazaei et al., 2017). This treatment produced OPCs at high-yield as indicated by robust Nkx-2.2 expression, and ultimately the OPC-specific antigen O4 (**Fig. 9*B***). We sought to generate more highly purified cultures of SMNs and OPCs for co-encapsulation in neural ribbons designed for the CNS (**Fig. 9*C***). Two additional purification steps were used prior to SMN-OPC co-culture and neural ribbon co-encapsulation that were MACS separation of OPCs with the O4 antigen (**Fig. 9*D***), and SMN-NS suspension culture. For encapsulation, OPCs were generated from hiPSCs expressing GFP. Enriched GFP-OPCs were retained and mixed 1:5 with SMNs and cultured for 1 w in modified N2B27 bundling medium (**Fig. 9*E***). At this point, SMN-OPC connected networks were encapsulated within 60 μm RGD-Col neural ribbons and embedded for culture and imaging using GFP and SMI312 immunofluorescence, shown at day 7 post-encapsulation (**Fig. 9*F***).

**Figure 9.**
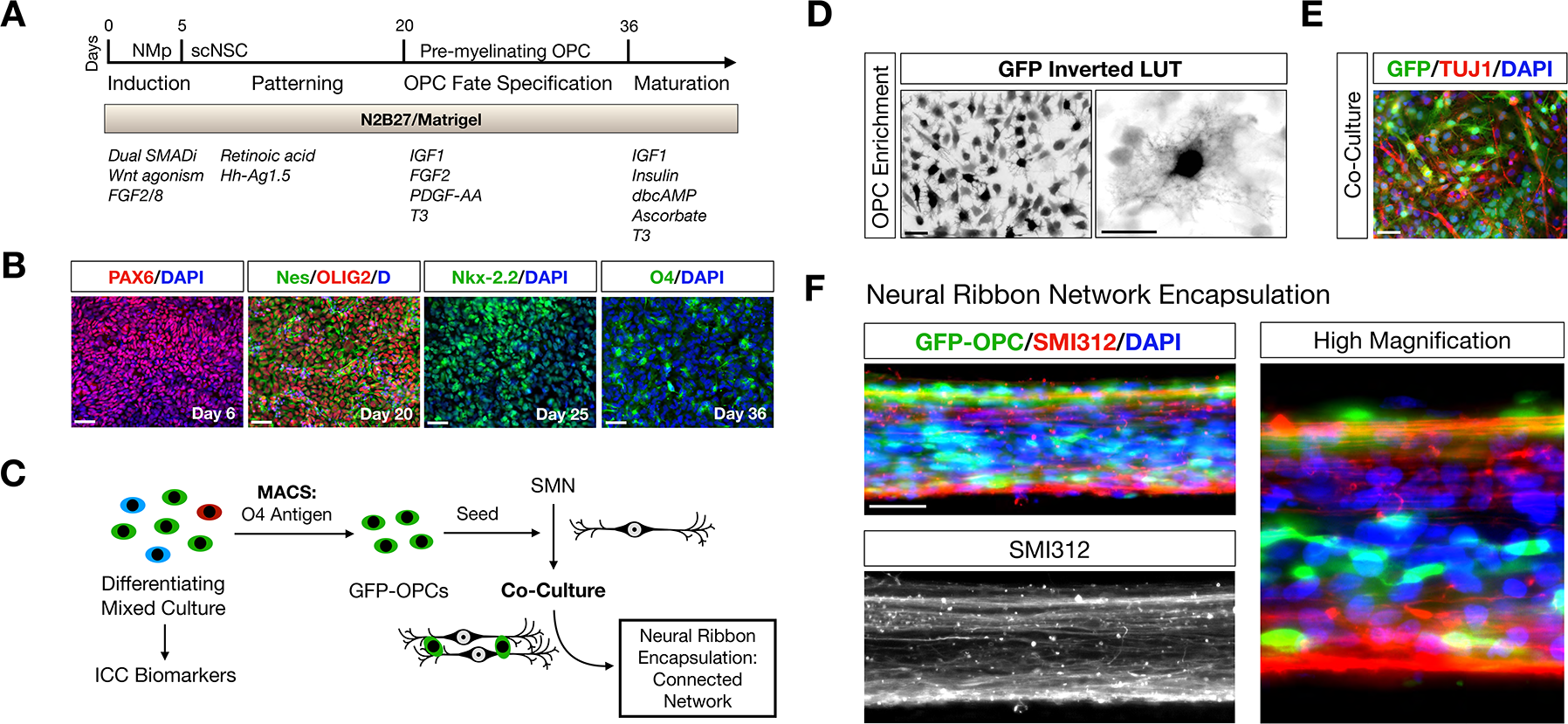
Encapsulating connected networks of spinal neurons and oligodendroglia. *A*, Overview of differentiation protocol used to generate oligodendrocyte progenitor cells (OPCs) through NMp and then multipotent scNSC stages. Small molecules and growth factors used in caudo-ventral patterning and maturation are listed. ***B***, IF images of OPC differentiation. Left-to-right: Pax6 (day 6); Nestin/OLIG2 (day 20); Nkx-2.2 (day 25); O4 (day 36). Cells counterstained with DAPI (D, nuclei). ***C***, GFP-OPC MACS separation scheme. GFP-OPCs were isolated from differentiating mixed cultures using magnetic beads functionalized with O4 antibody. Enriched GFP-OPCs were plated for further differentiation or seeded onto SMN cultures. ***D***, Adherent GFP-OPCs after MACS enrichment. ***E***, GFP-OPC co-culture with SMNs prior to encapsulation. ***F***, SMN-OPC connected networks were encapsulated within neural ribbons as a dual-component system. GFP and SMI312 immunostaining of neural ribbons is shown at day 7 post-encapsulation. High magnification image provided on right. Scale bars are 50 μm, and 10 μm in high magnification panel in (***D***).

### Neural ribbon linear placement and retention within collagen phantoms and the healthy rat spinal cord

We tested the ability to deliver and linearly place neural ribbons into 3D matrices *in vitro* and the healthy cervical spinal cord *in vivo* (**Fig. 10*A*** **and** **10*B***). To track neural ribbon constructs and cells, we labeled day 29 neuronal cultures with SPIO nanoparticles. Cultures were incubated overnight and neuron viability versus SPIO concentration was determined (**Fig. 11*A*** **and** **11*B***). Day 30 cultures labeled with ~100 μg Fe / 1 × 10^6^ cells (40 μl Miltenyi FeraTrack SPIO / 2 ml culture medium) were encapsulated with Fluorescein-Dextran (500 kDa) for visualization (**Fig. 11*C***), and shipped overnight to Houston Methodist. SPIO-Fluorescein neural ribbon segments (60 μm diameter, 3-4 mm length) containing day 32 neurons (~5,000 cells) were injected first into collagen phantoms and linearly placed as visualized by stereo microscopy (**Fig. 11*D***). To extend this technique to the rat CNS, we delivered individual SPIO-labeled day 32 neural ribbon segments to the healthy spinal cord (C4). The spinal column was imaged by ultra-high field micro MRI 24 h after transplantation (**Fig. 11*E*-11*G***). One sagittal plane (**Fig. 11*E***) and an axial series (**Fig. 11*F***) are provided as raw and processed images. Processed images display the T2 map computed from the gradient-echo signal (~T2* values) using a bone-range enhancer color map. In processed images, the implanted ribbon is visible in bright color. The so-called ribbon tip, neck and tail are visible in the axial series wherein the entry point of the neural ribbon tip occurs in the dorsal spinal cord and extends ~1.58 mm into ventral white matter. This positioning was further validated by measurements taken from the coronal plane (**Fig. 11*G***).

**Figure 10.**
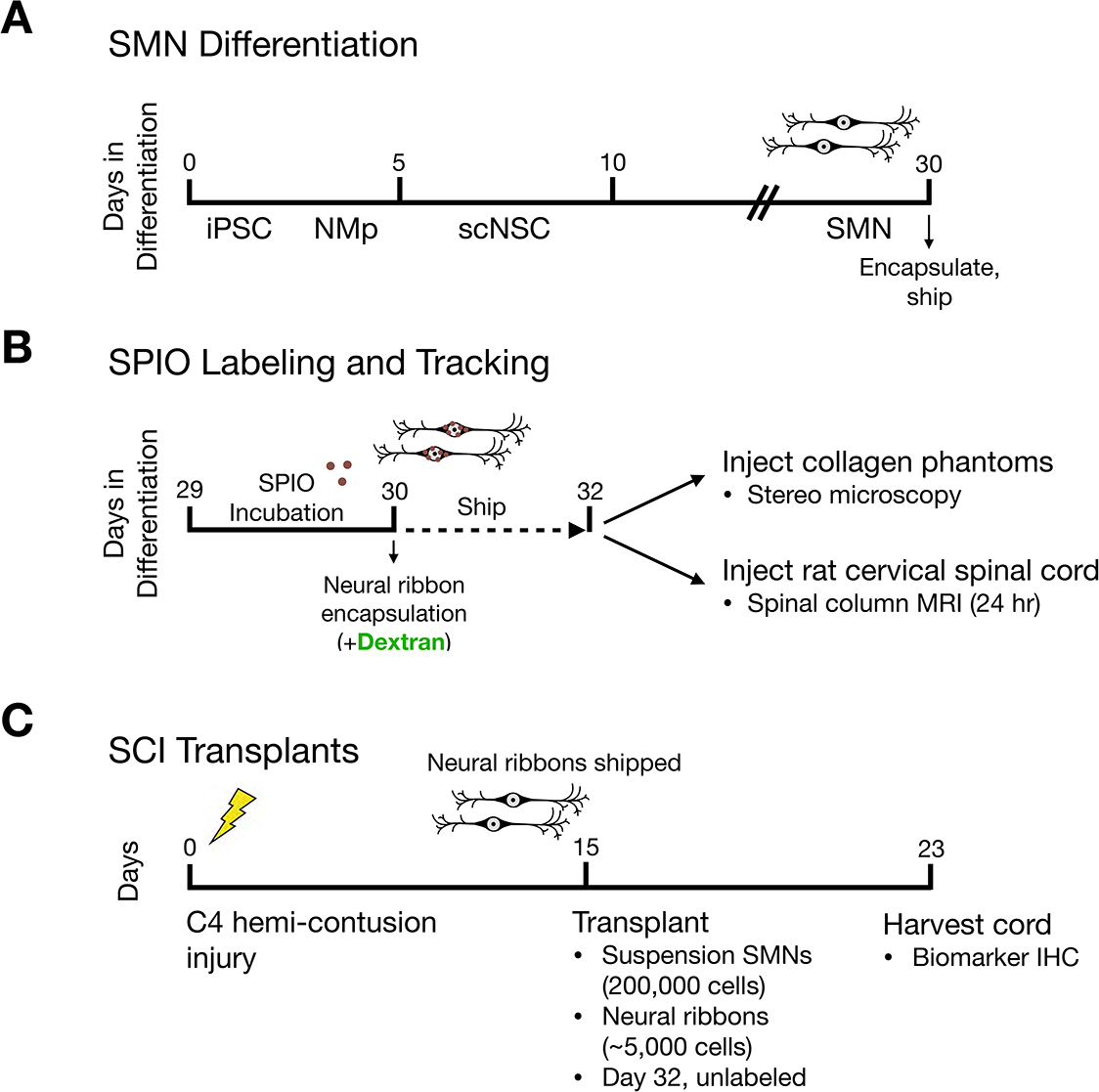
Experimental overview of *in vivo* investigations in rat. Experimental overview and timeline for pilot *in vivo* transplantation studies. ***A***, SMN differentiation, neural ribbon encapsulation, and shipping for rat *in vivo* delivery. ***B***, SPIO-labeling and tracking. SPIO-labeled cultures were encpasulated at day 30 and shipped overnight. At day 32, neural ribbons were injected into collagen phantoms and the healthy rat spinal cord for MRI. ***C***, C4 hemi-contusion injuries, transplant of neuronal suspensions (day 32 unlabeled, 200,000 cells derlivered) and neural ribbons (day 32 unlabeled, ~5,000 cells delivered) and downstream analysis.

**Figure 11.**
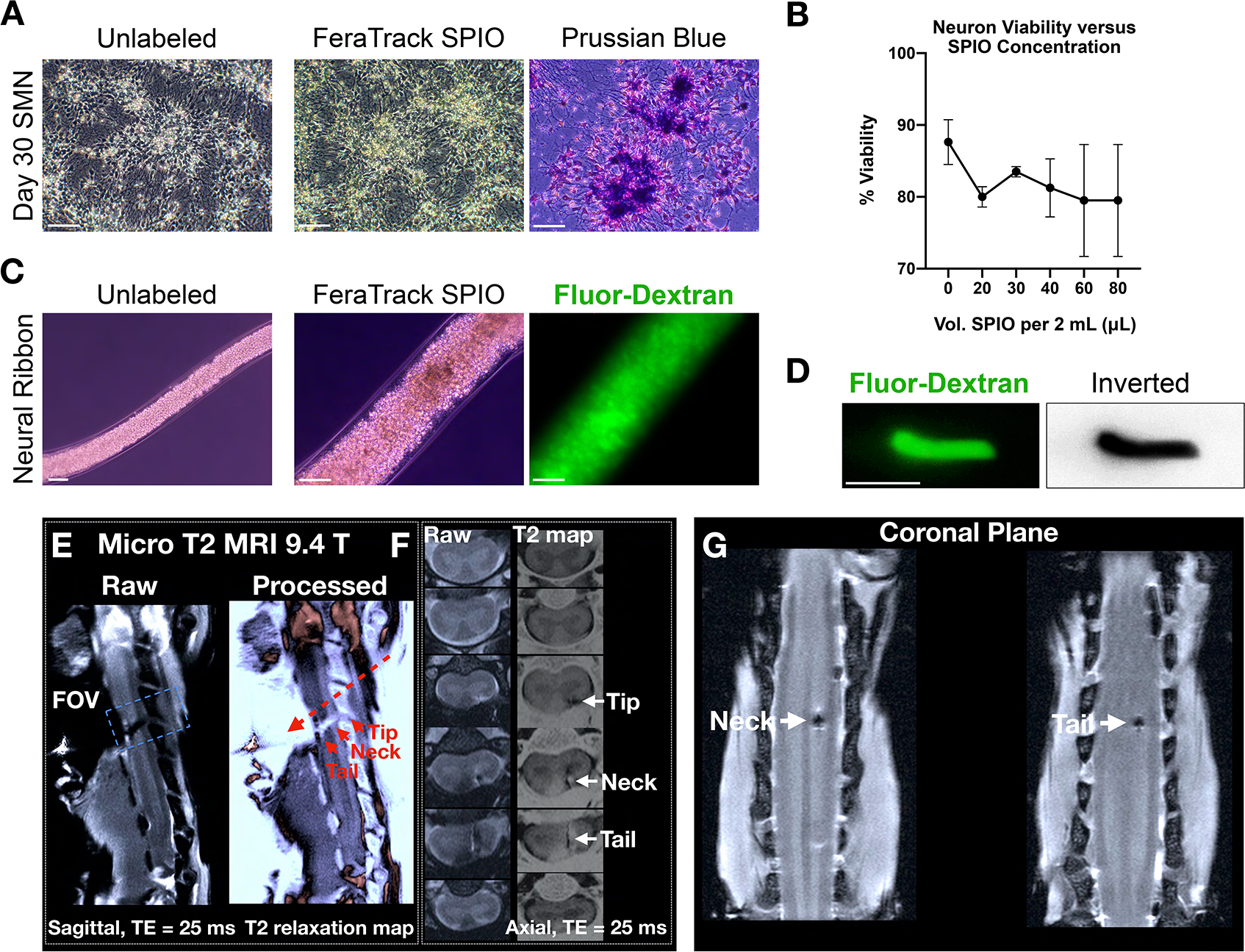
Delivery and linear placement of neural ribbons by injection *in vitro* and *in vivo*. ***A***, Unlabeled (left) and SPIO nanoparticle-labeled (middle) SMN cultures at day 30 validated with Prussian blue stain (right). ***B***, Neuron viability versus SPIO concentration after 24 h incubation. ***C***, Day 30 unlabeled (left; 10x magnification) and SPIO-labeled (middle; 20x magnification) SMN cultures encapsulated in neural ribbons with Fluorescein-Dextran (right). ***D***, ~4 mm SPIO-Fluorescein neural ribbon segment injected into collagen phantom. Inverted LUT is shown. ***E***, Sagittal plane raw (left, ribbon dark) and ~T2* processed (right, ribbon light) images depict neural ribbon transplant (tip, neck, tail regions; solid red arrows) and apparent direction of injection (dashed red arrow). ***F***, Axial planes of raw (left, ribbon dark) and processed (right, ribbon light) images from FOV in (***E***), highlighting ribbon tip, neck and tail. The neural ribbon tip is located at the dorsal position, while neck and tail regions extend across white matter for ~1.58 mm. ***G,*** Coronal plane views of neural ribbon neck and tail. Scale bars are 50 μm, and 3 mm in (***D***). Data represent mean ± s.e.m.

### Homotypic spinal neurons survive the contused injury cavity after both protected and unprotected delivery

We tested the ability to deliver and place viable hiPSC-derived cervical spinal neurons and neural ribbons into the contused spinal cord injury cavity (**Fig. 10*C*** **and** **Fig. 12**). Unlabeled neural cultures were encapsulated in 60 μm neural ribbons and shipped overnight from Albany, NY to Houston, TX. We used a well-characterized, reproducible rat model of cervical SCI (Mondello et al., 2015), wherein right-sided C4 hemi-contusion injuries were made using the electromagnetic spinal cord injury device (ESCID). 15 days post-injury, 8-weeks-old female Long-Evans rats (N = 4) received cervical intraspinal injections of single 3 - 4 mm neural ribbon segments (1 × 10^8^ cells/ml ribbon concentration, ~5,000 cell dose per animal). Using the neural ribbon platform, cell number is limited by geometric constrains of both the construct and the injury cavity size. For high dose suspension cultures, neural ribbon alginate was dissolved and cells were collected by centrifugation. ~200,000 cells were delivered per animal (N = 2). Eight days after transplantation (day 23 post-injury, day 40 from start of *in vitro* differentiation), spinal cords were harvested and processed for histological analysis (**Fig. 12**). Grafts were successfully delivered and retained in all six animals. At this time point, we observed robust survival of human spinal neurons (STEM121) delivered in suspension to within the injury cavity (**Fig. 12*A*** **and** **12*B***). Despite early glial scar formation (GFAP), this space-filling graft co-localized with SYN1 inside the cavity, and to some degree within adjacent host tissue (**Fig. 12*C***). Neuronal viability and STEM121/SYN1 co-localization in the cavity was preserved using smaller encapsulated grafts (40x reduction in cell number delivered), providing further support for neural ribbon-mediated neuroprotection (**Fig. 12*D*** **and** **12*E***). Neural network ribbons benefited robust graft penetration into host tissue by day 8 after transplantation (**Fig. 12*E***). Non-encapsulated grafts with low cell number (5,000 cells) delivered to the injury cavity have not been shown to survive (Olmsted et al., 2020).

**Figure 12.**
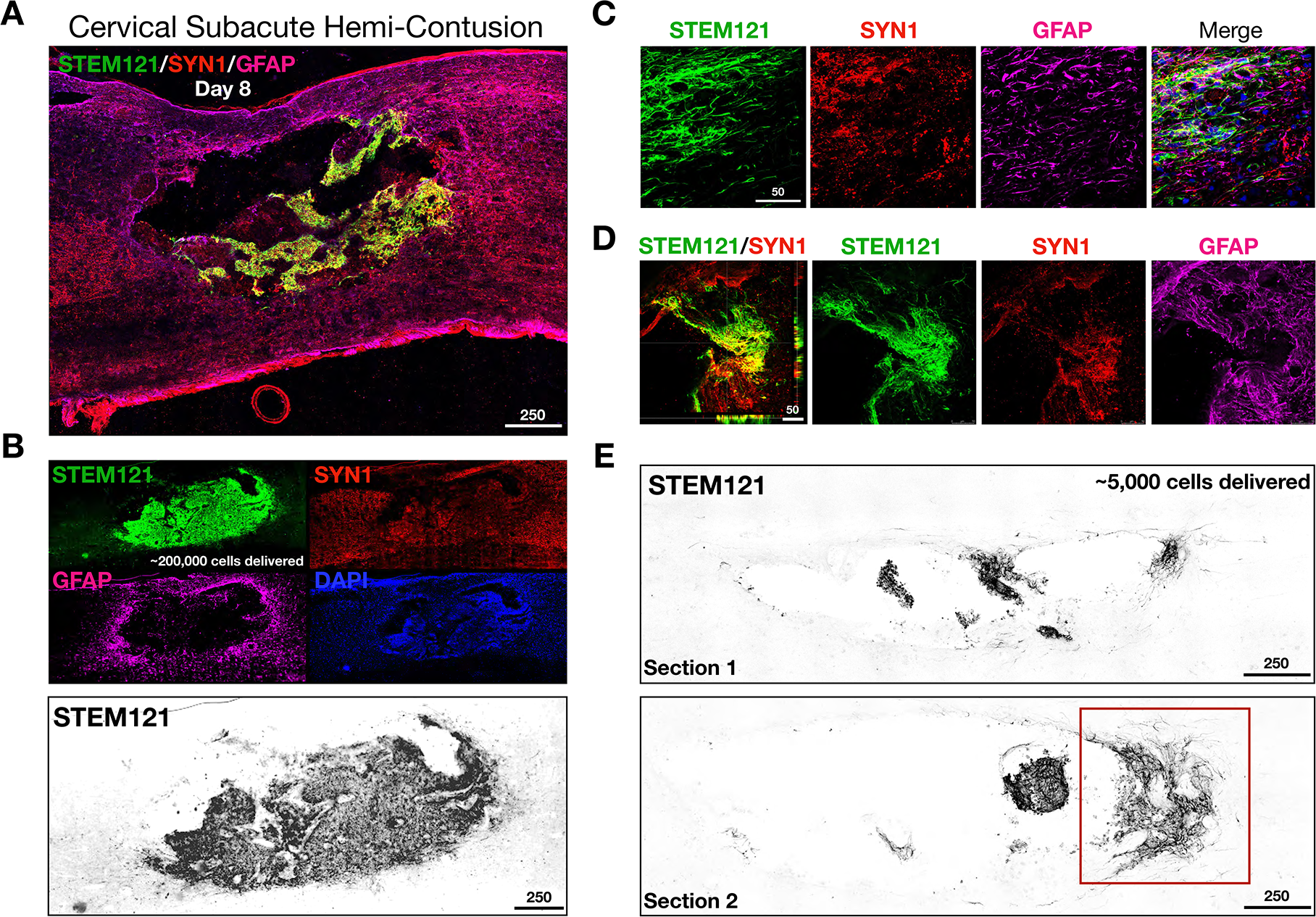
Survival and retention of encapsulated and non-encapsulated homotypic spinal neurons within the contused injury cavity *in vivo*. ***A***, Merged sagittal tile scan of C4 hemi-contusion injury grafted with ~200,000 hiPSC-derived unprotected spinal neurons (day 32 in differentiation) immunostained with STEM121 (human), GFAP, and SYN1 (top). ***B***, Central view of graft in (*A*) with individual biomarkers and STEM121 inverted LUT. ***C***, High magnification human neuronal projections into host parenchyma co-localize with SYN1. ***D***, Survival and engraftment of neural ribbon construct (~5,000 cells delivered) along the cavity border. STEM121 co-localization with SYN1 in the early glial scar 8 days after transplantation. ***E***, Two low magnification sagittal sections of neural ribbon graft from (*D*) shown with inverted LUT. Red box highlights neural network ribbon-supported penetration of graft into host tissue.

## Discussion

Neural circuits are comprised of populations of interconnected neurons and/or non-neuronal cell types that reliably output a specific response when stimulated. Here, we combine an advanced differentiation strategy through NMps for generation of SMNs and their anatomically relevant support cells into pre-formed immature neural networks as a novel, transplantable neural ribbon technology. These organized electrically-active networks provide an SMN regional match to advance anatomical domain-specific requirements to enable compatibility for microenvironment integration *in vivo*. As with other biomaterials studies, we also obtain neuroprotection benefits. Our approach is expected to benefit specialized treatment needs of neurological disorders that require stem cell replacement therapies that are adaptable to the degree of severity, neuroanatomical distribution, and the diversity of cell types impacted. For example, in Parkinson’s disease, advances in stem cell-based treatment are in part due to a unique understanding of the disease process in regards to etiology, neuroanatomy, and pathophysiology that centers around a single population of highly specialized dopaminergic neurons (Barker et al., 2017). In the SCI field, traumatic injury introduces increasing degrees of complexity, as therapeutic interventions must consider the neuroanatomical gamut of rostral-caudal and dorsal-ventral axes, with the possibility of impacting numerous neural circuits and cell types. The future clinical scenario must also address the complex pathophysiological response to injury over time, as well as the degree of a refractory microenvironment. The current primary method for therapeutic grafting in the spinal cord is by delivery of high-dose dissociated NSCs in suspension, with the expectation that a substantial but unpredictable percentage of these cells will not survive. Despite demonstration of safety in an initial phase 1 human clinical trial (Curtis et al., 2018) and achievement of graft long-distance axonal projections in pre-clinical animal models (Lu et al., 2014; Brock et al., 2018), delivery of unprotected NSCs has proved insufficient for restoring complex cellular organizations to pre-injury states. Additionally, the large cell loss of transplanted cells creates a challenge in regard to predictable outcomes if more complex multi-cell type suspensions are used. The ability to successfully deliver and retain cells at the injury site is therefore lagging behind our advancements in the understanding of normal and disease states following injury. Reproducible methodologies for delivery of pre-configured functional networks with regionally-defined cellular components that enable neuroprotection, integration, and patient-specific adaptations are therefore of high priority.

By developing encapsulation technology for maturing neurons and neural co-cultures that supports and stabilizes connectivity for early network formation we can begin to address this need. Importantly, this neural ribbon technology enables the delivery of functional homotypic neurons to the CNS that is in contrast to the vast majority of stem cell transplantation studies centered around multipotent NSCs. In particular, we generated hiPSC-derived cervical spinal neurons (Davis-Dusenbery et al., 2014; Sances et al., 2016; Trawczysnki et al., 2019) using developmentally relevant NMp intermediates that have been described (Gouti et al., 2014; Lippmann et al., 2015). These cells were appropriately patterned to express tissue specific-targets for expediting spinal cord engraftment. The importance of homotypic patterning (Dulin et al., 2018) is underscored by our observations of survival and neurite extension of unprotected spinal neurons delivered in suspension to the injury cavity (~200,000 cells). As neural ribbons, the neuroprotective role of encapsulation is supported by survival of grafts containing only ~5000 cells, as well as their ability to rapidly penetrate into the host parenchyma.

The requirement for 3D methodologies to accelerate neuronal maturation in terms of excitability, input integration, spontaneous firing properties, connectivity, and network formation has been emphasized in several recent studies (Bakooshli et al., 2019; Izsak et al., 2019; Faustino Martins et al., 2020). Two of these studies further model maturation of neuronal interactions with non-neuronal cell types in a 3D context, particularly with skeletal muscle, in self-organizing trunk organoids (Faustino Martins et al., 2020) and advanced neuromuscular co-cultures (Bakooshli et al., 2019). A study by Izsak et al. (2019) employed microelectrode array (MEA) technology to demonstrate rapid achievement of population bursting properties by seeding 3D neural aggregates onto arrays versus dissociated cells (Izsak et al., 2019). Accordingly, we observe glutamate-responsive network bursting events by incorporation of SMNs into neural ribbons that were not detected in adherent cultures. Network bursts were retained in neural aggregates even after the dissolution of neural ribbon hydrogel. In summary, we encapsulate transplantable populations of neurons and additional neural cell types interconnected by synapses that activate when stimulated by external cues. Similar to organoids, which do not currently extend directly to transplantation into the spinal cord, neural ribbons are expected to provide a novel tool to study the early formation of neuronal networks and ultimately the development and physiology of high-fidelity neural circuits. Accordingly, neural ribbon constructs may be useful *in vitro* for basic neuroscience investigations and drug discovery. This technology constitutes an important first step towards greater control over direction of functional neuronal networks in neuroprotective platforms that support rapid engraftment and survival in the CNS at controlled cell dose.

## Acknowledgements

This work was funded by the New York State Department of Health (NYS DOH) Spinal Cord Injury Research Board (NYSCIRB), Projects to Accelerate Research Translation (PART) award C33278GG and SUNY Polytechnic SEED award 917035-21 and used a published line developed through previous New York State stem cell research (NYSTEM) funding. RNA-Seq was performed at the SUNY Buffalo Genomics and Bioinformatics Core. This work was also supported by NSF IOS1655365 and SUNY Albany Research Foundation grants to A.S.

## Notes

**Competing financial interests:** The authors declare no competing financial interests.

### Competing Interest Statement

The authors have declared no competing interest.

